# The structure of the monobactam-producing thioesterase domain of SulM forms a unique complex with the upstream carrier protein domain

**DOI:** 10.1101/2024.04.06.588331

**Authors:** Ketan D. Patel, Ryan A. Oliver, Michael S. Lichstrahl, Rongfeng Li, Craig A. Townsend, Andrew M. Gulick

**Author notes:** Address correspondence to Andrew M. Gulick, The Jacobs School of Medicine & Biomedical Sciences, 955 Main Street, Buffalo, NY, 14203.

## Abstract

Nonribosomal peptide synthetases (NRPSs) are responsible for the production of important biologically active peptides. The large, multidomain NRPSs operate through an assembly line strategy in which the growing peptide is tethered to carrier domains that deliver the intermediates to neighboring catalytic domains. While most NRPS domains catalyze standard chemistry of amino acid activation, peptide bond formation and product release, some canonical NRPS catalytic domains promote unexpected chemistry. The paradigm monobactam antibiotic sulfazecin is produced through the activity of a terminal thioesterase domain that catalyzes an unusual β-lactam forming reaction in which the nitrogen of the C-terminal *N*-sulfo-2,3-diaminopropionate residue attacks its thioester tether to release the β-lactam product. We have determined the structure of the thioesterase domain as both a free-standing domain and a didomain complex with the upstream *holo* peptidyl-carrier domain. The structure illustrates a constrained active site that orients the substrate properly for β-lactam formation. In this regard, the structure is similar to the β-lactone forming thioesterase domain responsible for the production of obafluorin. Analysis of the structure identifies features that are responsible for this four-membered ring closure and enable bioinformatic analysis to identify additional, uncharacterized β-lactam-forming biosynthetic gene clusters by genome mining.

---

Many microbes produce a variety of natural products that are secreted from the cell where they play roles in adaptation to numerous changing environments. These small molecules may facilitate competition with other organisms or play a role in the acquisition of important nutrients. An important class of natural products are peptide and peptide-like molecules produced by the nonribosomal peptide synthetases (NRPSs). Not constrained by the synthetic limitations of the ribosome and amino acyl-tRNA synthetases, the NRPSs form products from hundreds of unique building blocks (1,2). The diversity of the peptide products is enhanced not only by the wide spectrum of substrates but also by additional changes to cross-link, cyclize, and otherwise modify these molecules during their biosynthesis. NRPS products range in size from small cyclic dipeptides to large macrocycles containing more than twenty amino acids. Much effort has gone into the identification of NRPS biosynthetic gene clusters and the molecules that are produced by the encoded proteins, with multiple informatic tools available for their identification, analysis, and classification (3-5).

The NRPSs are large, modular proteins that function as assembly lines for the stepwise construction and extension of a peptide product. Each module is composed of a defined set of catalytic domains, which is generally responsible for the incorporation of a single amino acid building block. Each module houses a peptidyl carrier-protein domain (PCP) that contains a conserved serine residue that is post-translationally modified with a molecule of phosphopantetheine. The pantetheine thiol serves as the binding site for the amino acid substrate and the growing peptide. In addition to this carrier domain, NRPS modules generally contain adenylation domains that are responsible for recognizing and loading the substrate onto the PCP and condensation domains that catalyze peptide bond formation simultaneously transferring the amino acid or upstream peptide to a downstream PCP domain, thereby extending the peptide by one residue. While the covalent tethering of the nascent peptide to the PCP facilitates coordination of the catalytic steps, this feature also requires a final catalytic step to release the peptide from the terminal PCP domain. This step is most commonly carried out by a thioesterase domain that catalyzes hydrolysis of the peptide thioester with the pantetheine cofactor or, often, a macrocyclization step carried out when a nucleophilic group on the peptide attacks the thioester (6,7). Cyclization can occur through the N-terminal amine of the peptide or internal side chain groups from lysine, serine, threonine or other nucleophiles.

The thioesterase domain belongs to a large α/β hydrolase superfamily that catalyze hydrolysis of multiple substrates (8,9). These enzymes contain a conserved fold of a central 7-or 8-stranded β-sheet with surrounding α-helices. Between strands β6 and β7 is a long loop generally containing two helices that form a lid over the active site. The catalytic machinery of the thioesterase domain contains a nucleophilic serine or less common cysteine residue that is activated by an aspartate and histidine dyad that form a hydrogen bonding network with the nucleophile (6,9). The reaction mechanism employs two steps to release the peptide from the terminal PCP domain. First, the serine or cysteine attacks the thioester to transfer the peptide from the PCP and form an acyl-enzyme intermediate. This intermediate is resolved through a second off-loading step that is specific to the cyclization or hydrolysis necessary for each enzyme.

Previously, crystal structures have been determined for eight NRPS thioesterase domains. Unlike other NRPS catalytic domains, for which numerous structures have been solved complexed with informative ligands or paired with the carrier protein domains (10-12), the structural foundation of thioesterase domain function is much less well-characterized. There are only two structures of thioesterase domains that provide insight into their interactions with substrates. The thioesterase domains from the nocardicin and valinomycin pathways have been crystallized with ligands bound to the nucleophile to provide insight into the transiently formed acyl-enzyme intermediate. In the case of the thioesterase domain from the nocardicin protein NocB, a phosphonate analog of the peptide substrate was designed that reacted with the catalytic serine to identify the location of the terminal three residues of the peptide just prior to release (13). In the valinomycin producing Vlm2 enzyme, a protein engineering approach allowed the replacement of the catalytic residue with 2,4 diaminopropionate that reacted stably with the peptide (14). Finally, only a single thioesterase domain, present in the enterobactin NRPS EntF (15,16), has been characterized as a complex with the PCP domain.

Among the most interesting and important NRPS products are peptide antibiotics that range in size as large as the glycopeptide and lipopeptide antibiotics containing ten or more amino acids (17,18). Among the antibiotics produced by NRPS pathways are β-lactam antibiotics, such as penicillin, cephalosporin, and the monocyclic nocardicin and sulfazecin (19). Interestingly, the β-lactam ring is produced in these different molecules through different enzyme activities. A non-heme iron-dependent oxygenase catalyzes the production of the penicillins and cephalosporins from an NRPS tripeptide precursor that has been released from the enzyme (20). Other systems integrate the β-lactam producing step directly into the NRPS, with ring formation occurring in the nocardicin pathway from an integrated condensation domain (21), while the activity of a thioesterase domain catalyzes the cyclization in sulfazecin biosynthesis (22,23). These strategies in NRPS systems contrast with an ATP dependent β-lactam synthetases that catalyze the ring closure in the production of the β-lactamase inhibitor clavulanic acid (24) and the broad-spectrum carbapenems (25,26) by NRPS-independent pathways (27,28).

Sulfazecin and its stereoisomer isosulfazecin were first identified in the early 1980s from a producer strain of *Pseudomonas acidophila* (29,30). Sulfazecin consists of a tripeptide derivative formed from γ-glutamate, D-alanine, and an azetidin-2-one ring containing a sulfonate linkage to the nitrogen and a methoxy group added to the lactam ring. Sulfazecin binds to the *E. coli* penicillin binding proteins and was bacteriostatic at 6.25 µg ml^-1^. Isosulfazecin, the L-stereoisomer at the alanine residue was reported to have slightly weaker antibacterial activity against some strains (31).

The gene cluster was identified through transposon mutagenesis and screening for the inability to inhibit *E. coli* growth. A single mutant strain was isolated from the producer *P. acidophila*, allowing the discovery of the biosynthetic cluster (22). We note that the producing organism has recently been reclassified and is recognized in NCBI as *Paraburkholderia acidicola* (32,33).

The biosynthesis of sulfazecin (Figure 1) employs a biosynthetic gene cluster containing two NRPS proteins, SulI (1089 residues) contains the initiation module that incorporates the γ-glutamate residues, while SulM (2984 residues) contains the second and third module (22). Also found within the cluster are SulG and SulH, involved in the production of L-2,3-diaminopropionate (L-DAP), and SulN, a sulfotransferase, and SulO and SulP, which were shown to install the methoxy group (23). The presence of SulG and SulH suggested that the β-lactam ring in sulfazecin might be derived from cyclization of L-DAP. Biochemical analysis confirmed the incorporation of L-Ala and L-DAP by the adenylation domains of the SulM modules. Subsequently, it was shown that the β-lactam ring of sulfazecin was formed by the activity of the thioesterase domain, which catalyzes the intramolecular attack of the β-amine on the thioester following *in trans* SulN-catalyzed *N-*sulfonation (23). Following release of the desmethoxy sulfazecin, the methoxy group is installed in two steps by the SulO and SulP enzymes.

**Figure 1.**
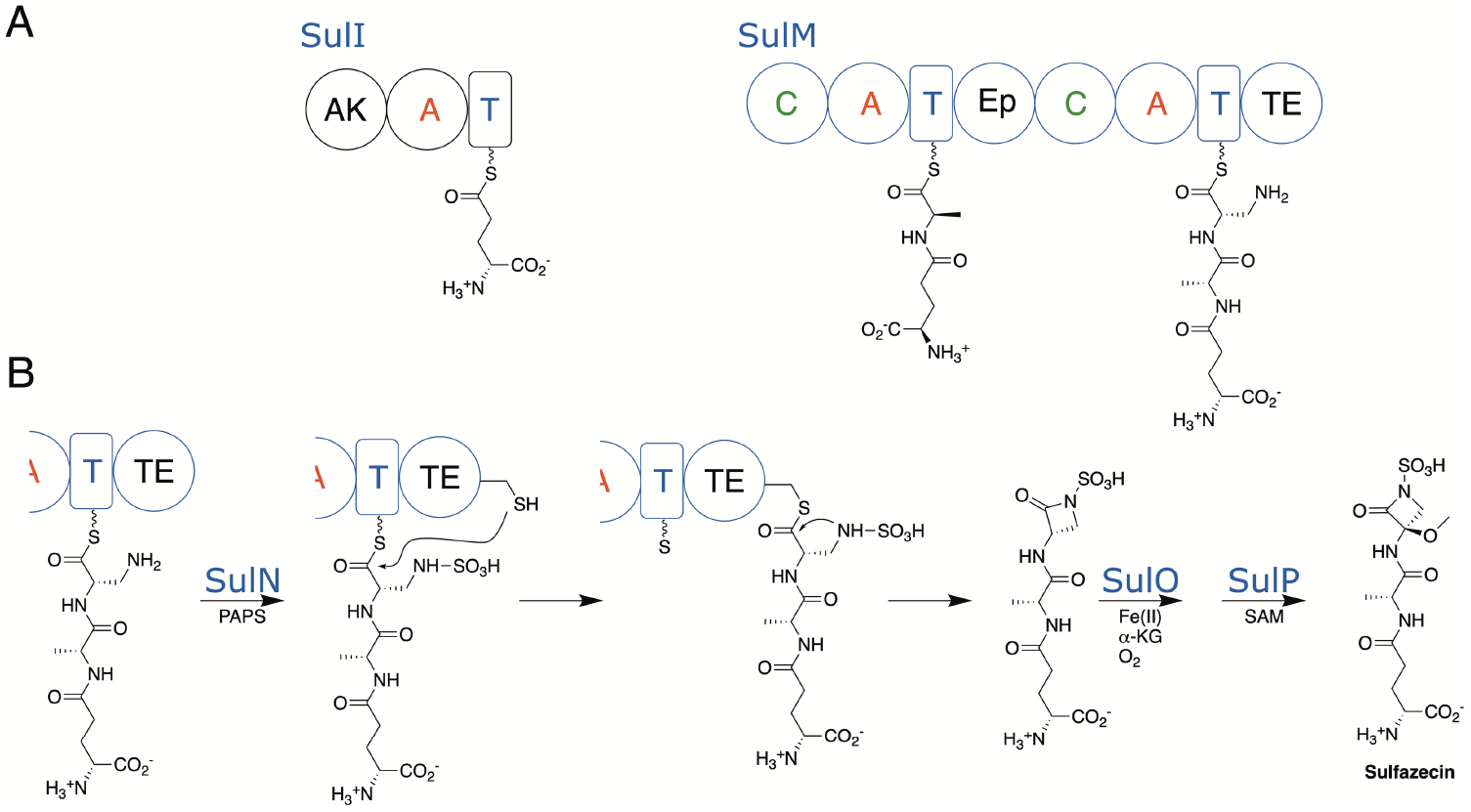
Biosynthesis of Sulfazecin. A. Two NRPS proteins, SulI and SulM, harbor three modules necessary to produce the desmethoxysulfazecin. The SulI module contains an adenylylsulfate kinase, responsible for production of PAPS, as well as the initiation module for D-glutamate. SulM contains two modules that incorporate L-alanine, which is epimerized to the D-stereoisomer, in module 2 and L-DAP in module 3. B. The final steps in sulfazecin production include the *in trans* sulfonation catalyzed by SulN, and then β-lactam formation by the thioesterase domain, resulting in the production of desmethoxysulfazecin. Two final steps, catalyzed by the nonheme iron oxygenase SulO and methyltransferase SulP produce sulfazecin.

The terminal β-lactam forming catalytic activity of SulM TE domain motivated us to solve the structure of this domain. Here we present the structure of the thioesterase domain on its own and as a complex with the upstream *holo*-PCP domain. The structure provides a view of the functional interaction between the carrier and the thioesterase domain, which illustrates a distinct interaction compared to EntF, the only prior structure of a PCP-thioesterase interaction. Analysis of the active site suggests that the thioesterase domains of β-lactam and β-lactone forming enzymes provide a constrained architecture for the nucleophilic group on the Cβ amine or hydroxyl of the C-terminal amino acid of the peptide to attack the thioester linkage for formation of the four-membered ring product.

## Results and Discussion

### Structure of the SulM TE domain

There are currently structures for eight thioesterase domains (12) from modular NRPSs solved as single domain recombinant proteins or as part of a larger multidomain construct. The proteins represent distantly related enzymes, with amino acid pairwise sequence identities that range from 17 to 28 % between different members (Figure S1). The structures include the single domains from surfactin (34), fengycin (35), valinomycin (14), nocardicin (13), and skyllamycin (36) NRPS termination modules. Additionally, there are several structures of the thioesterase domains from larger multidomain proteins including a PCP-TE complex from EntF (15,16), which shows the interaction of the PCP and TE domains. The other NRPS thioesterase domains, involved in the production of surfactin (37), enterobactin (38), and obafluorin (39), as well as an uncharacterized NRPS from *A. baumannii* (40), illustrate the position of the thioesterase domain relative to a complete termination module. In these structures, the thioesterase domain is highly dynamic and adopts multiple positions relative to the core of the module and does not interact functionally with the upstream PCP domains.

To explore the structural foundation for β-lactam formation by the TE domain of SulM in sulfazecin biosynthesis, we determined the structure of the domain in the presence and absence of the partner carrier protein. Protein constructs from five cut sites upstream of the TE coding sequence were selected for expression tests. The structure of the optimized genetically truncated thioesterase domain, which we refer to as SulTE, was determined (Figure 2). The protein has a typical α/β hydrolase fold with seven central β strands (β2 to β8) flanked by four α helices on one side and three helices on another side (Figure S2A). The first strand of the canonical α/β hydrolase fold is missing and hence the structure starts with the second β-strand, which is anti-parallel to the remaining six strands. A very long sixth α-helix along with seventh helix and the following loop (residues 2860 to 2877) form the lid region.

**Figure 2.**
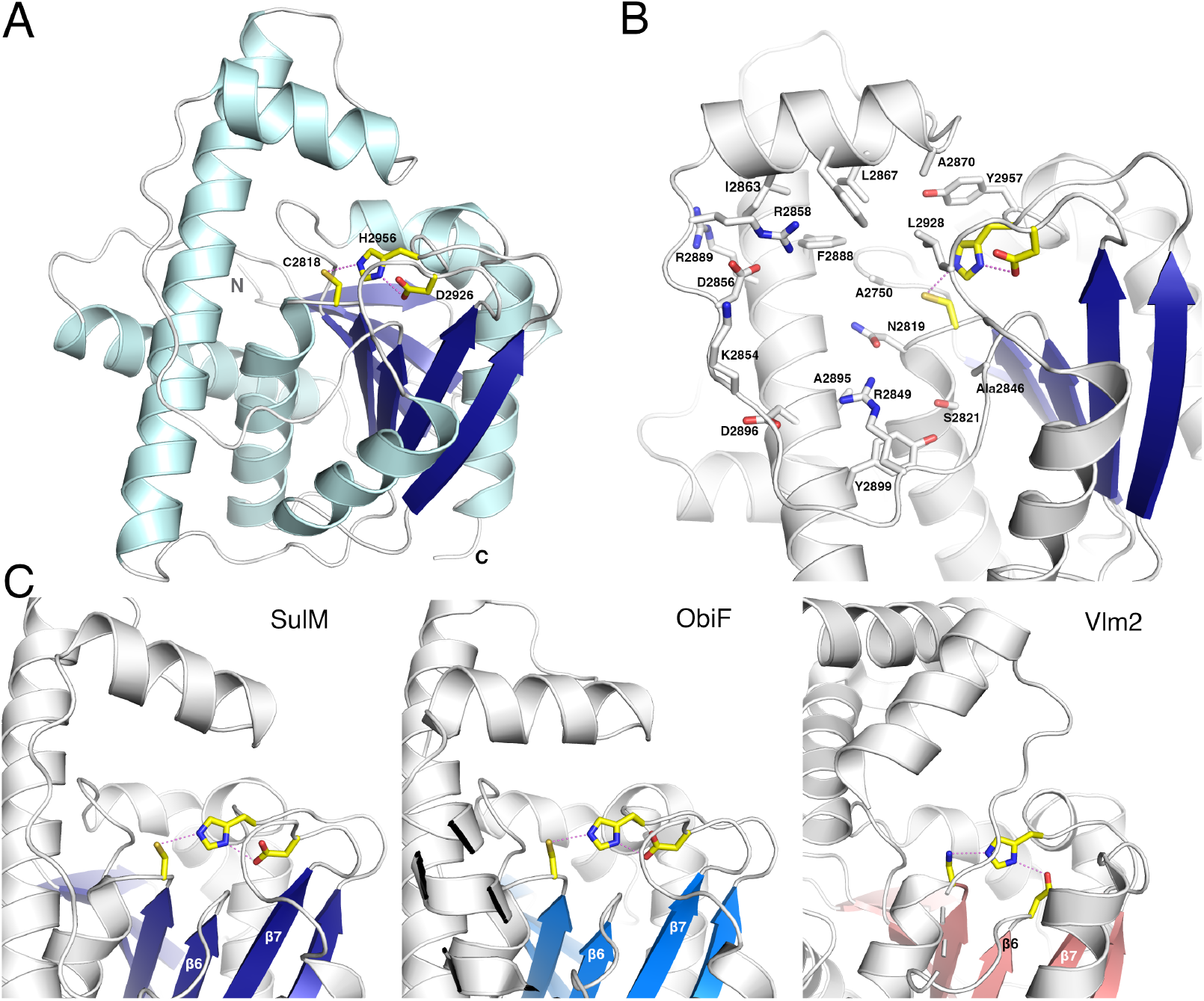
Structure of SulM thioesterase domain. The SulM thioesterase domain adopts a conventional α/β hydrolase fold with a central β-sheet surrounded on both sides by α-helices. A. The catalytic triad illustrates a nucleophilic cysteine residue Cys2818 that is positioned on the loop following strand β5, interacting with His2956 and Asp2926. B. The active site pocket is formed from residues on the lid loops and lid helices. C. SulM and ObiF thioesterase domains contain a catalytic triad in which the aspartic acid is positioned on the loop following strand β7. In contrast, the aspartic acid residue of Vlm2, like all other structurally characterized NRPS thioesterase domains, is positioned in the loop following strand β6.

A comparison of the SulM thioesterase domain with earlier NRPS crystal structures shows that the protein adopts the conventional overall hydrolase fold. The root mean square (rms) displacement of pairwise comparisons of SulM with the remaining proteins ranges from 1.3 to 3.2 Å (Figure S1), excluding the dynamic lid loops, with most pairwise comparisons showing rms displacements less than 2Å.

The active site residues of the SulM thioesterase domain, Cys2818, Asp2926, and His2956, form the catalytic triad where the central histidine is 3.2 Å and 2.7 Å apart from the cysteine and aspartic acid residues, respectively. The nucleophile Cys2818 is present at the turn following the β5 strand, while Asp2926 is located on the loop that follows the β7 strand and His2956 is positioned on the loop after β8 strand.

This organization of the catalytic triad is distinct from most NRPS thioesterase domains. Unlike the position of the aspartic acid residue on β6 (position I) seen in other NRPS TE domains, only ObiF1 (39) and SulTE reposition the aspartic acid residue to the loop following strand β7 referred as position II (Figure 2A). This arrangement has also been observed in some type II proof-reading thioesterases such as RifR and RedJ (41,42) and, indeed, is the more common configuration in other members of the α/β hydrolase superfamily (8,9). To explore the impact of this altered positioning, the aspartic acid residue of ObiF1 was mutated to an alanine, resulting in a nearly 10-fold reduction in production formation (39). In ObiF, relocation of the aspartic acid to the location following strand β6 with a double mutant failed to restore activity, leading to the hypothesis that the unusual triad orientation seen in ObiF and presumably SulM is necessary to prevent steric hindrance of the loaded acyl enzyme intermediate.

We compared the nature of the specific reaction chemistry (hydrolysis, cyclization, or β-lactam/lactone formation), the nature of the nucleophile (serine or cysteine), as well as the peptide substrate for the different NRPS enzymes to correlate with the unusual catalytic triad of the sulfazecin and obafluorin proteins. The thioesterase domains of both ObiF and SulM contain a cysteine as the catalytic nucleophile and both require the attack of the β-carbon hydroxyl or amine on the thioester to form the lactone or lactam ring respectively. Thus, the relocation of the catalytic aspartic acid may facilitate the necessary configuration of the terminal residue of the peptide when bound as an acyl-enzyme intermediate to the nucleophilic cysteine to properly allow the β-lactam or β-lactone forming reaction to proceed. Of the other structurally characterized proteins, only the uncharacterized AB3403 from *A. baumannii* also contains a nucleophilic cysteine, although it contains an aspartic acid residue of the triad that follows the β6 like other NRPS proteins (position I).

The other feature that stands out among the characterized NRPS systems is the size of the peptide product that will be situated on the catalytic nucleophile as part of the catalytic mechanism. The ultimate structures of the products released from these thioesterase domains (Figure S3) highlight that the sulfazecin and obafluorin molecules are much smaller (mw < 370 Da) than the products of the other NRPS systems (mw ranging from 670 to 1470 Da). Thus, it is possible that the positioning of the catalytic triad in these NRPS enzymes with larger peptide products may be necessary to provide sufficient space in the active site to accommodate the large peptide.

### The lid helices of SulTE highlight structural diversity among the NRPS thioesterase domains

Alignment of the structurally characterized NRPS thioesterase domains illustrates that, despite the low sequence homology, the main secondary structural elements of the α/β superfamily superimpose for all family members. The distinguishing feature, however, in comparison of the structures is the region known as the lid loop. Thioesterase domains of the α/β superfamily generally contain two helices located between strands β6 and β7 (6). These two helices are positioned near the active site and interact with the loaded substrate in the acyl-enzyme complexes of NocB or Vlm2 (13,14). We examined the NRPS thioesterase domain structures and noted remarkable diversity in the length and helical content of this lid region (Figure S2B). The lid loop joining β6 and β7 ranges in size from 54 to 101 residues, containing as few as two and as many as five α-helices (Table S1).

The SulM thioesterse domain lid loop, which spans residues Gly2844 to Val2918, contains two helices. The second of these is the longest helix of a characterized NRPS thioesterase domain. This helix in the SulM thioesterase aligns with the final helix of all NRPS thioesterase structures. The SulM lid contains an extension of one additional turn of the helix compared to closest structural homolog ObiF and is longer by two to four helical turns than the other structures. The N-terminus of the helices align; however, they differ in the length with longer helices projecting away from the active site.

A second loop, joining strands β7 and β8, also adopts very different configurations in the different structures although it does not interact with ligands in the NocB or Vlm2 thioesterase domain structures (Figure S2). While not as dramatic as the lid region, the loop joining β7 and β8 ranges from 19 to 26 residues in length, generally containing a single α-helix of 5 to 16 residues (Table S1). This second, variable loop contains no α-helices in the SrfA-C structure (34,37), while the same region in the fengycin thioesterase domain (35) contains a split helix that is interrupted by a residue that adopts a left-handed helical conformation.

In the SulTE structure, the loop that follows the β7 strand runs from Gly2923 to Val2946 and contains an unusually long α-3_10_ composite helix. The first two residues of this helix, Tyr2932 and Pro2933, adopt a traditional α-helical conformation while the following nine residues form a 3_10_ helix. Among known NRPS thioesterase domain structures, only SulM and ObiF1 show α-3_10_ composite helix after β7 strand. As this loop contains the catalytic aspartic acid residue in these two proteins, the configuration of this loop may relate to the need to position this residue closer to the active site. In contrast, the loop after the β7 strand is further away from the active site in enzymes having the acidic aspartate residue after β6 strand (position I).

### Structure of the SulM TE domain in complex with the upstream PCP

We determined the didomain PCP-TE of SulM, in which the PCP was in the *holo* state loaded with the pantetheine cofactor and the catalytic cysteine of the thioesterase domain was mutated to an alanine. We designed a peptide mimic of the sulfazecin tripeptide by loading the pantetheine with the γ-D-Glu-D-Ala-L-Glu, where the C-terminal glutamate potentially forms anionic interactions of the natural *N-* sulfonated DAP substrate (Figure 3). This non-reactive substrate surrogate was easily prepared analogously to the native γ-D-Glu-D-Ala-L-Dap sulfamate (23). The PCP-TE structure was determined and refined to a resolution of 2.73 Å resolution. The structure contained two independent chains in the asymmetric unit, each containing residues Glu2660 through Cys2979. The N-terminus of the protein also contained four residues from the purification tag, while the C-terminal five residues were disordered. The two chains superimposed with a rms displacement of 0.4 Å over the full length of both domains, indicating that the orientation of the PCP relative to the thioesterase domain was the same in both chains. Contiguous electron density the pantetheine was observed on Ser2688 of the PCP domains in both chains (Figure S4).

**Figure 3.**
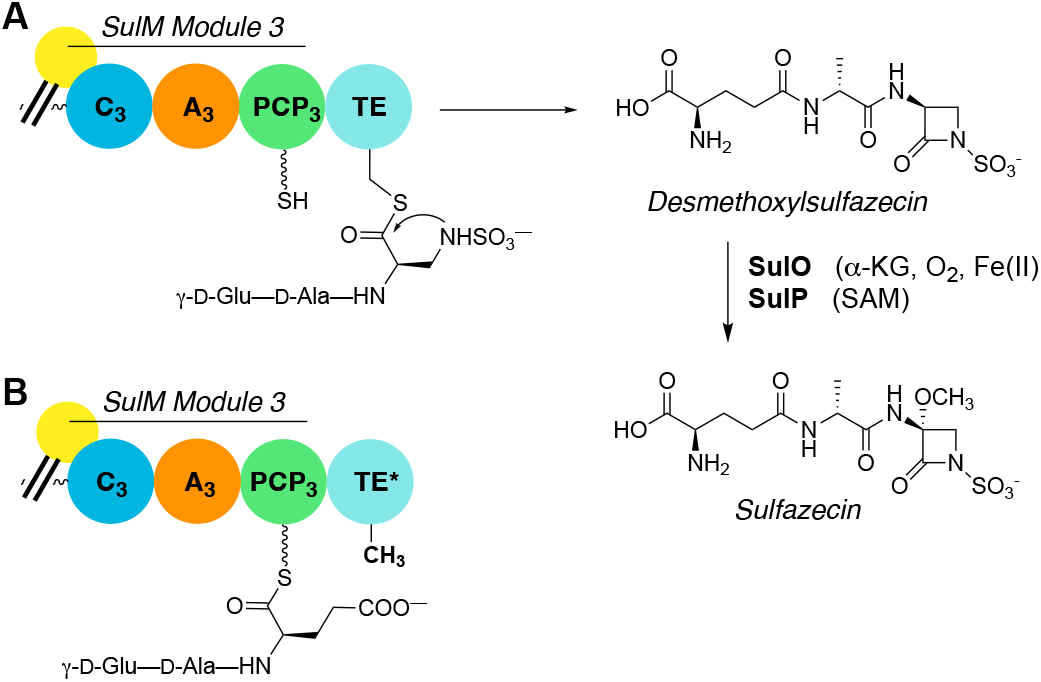
Examination of monobactam formation in SulTE. A. γ-D-Glu-D-Ala-L-Dap linked to the panethiene arm of PCP_3_ in the NRPS SulM is sulfonated *in trans* to the corresponding sulfamate and delivered to the SulTE catalytic Cys2818 by transthioesterification and cyclized to the monocyclic β-lactam product and released. Subsequent oxidation by SulO and *O*-methylatioin by SulP yield the monobactam sulfazecin. (B) The *C*-terminal amino acid of the tripeptide in A was replaced with L-Glu to mimic the distal charge of the native sulfamate.

The conserved position of the pantetheine attachment at the start of helix α2 of the PCP directs one face of the carrier domain towards the neighboring catalytic domains, with residues from α2, α3, and the loop joining helix α1 and α2 playing primary roles in the interactions (11). Nonetheless, there are subtle differences in the residues/interfaces that form the interaction and in the trajectory of pantetheine arms relative to the PCP domains when interacting with the alternate domains (Figure S5). Although confirmed to be present in the protein prior to crystallization set up, no density was present to warrant inclusion of the tripeptide mimic in the final model. As such, the pantetheine is modeled as a free thiol. Whether the peptide was hydrolyzed over the course of the crystallization experiment was not investigated.

Comparison of the structure of the isolated thioesterase domain with the didomain structure showed small conformational changes in the lid helices. The rms displacement of the Cα positions in the TE domain is 0.5 Å (over 251 Cα atoms). All secondary structure elements superimpose well, with only the loop joining the αL1 and αL2 adopting a slightly different position in the didomain structure to increase the size of the pantetheine binding tunnel (Figure 4). The pantetheine reaches from the PCP domain into the active site, positioning the sulfhydryl 4 Å from Cβ of the alanine residue that replaced Ser2818. The pantoic acid dimethyl group of PPant interacts similarly in a hydrophobic pocket formed by Leu2874, Phe2884. The pantetheine hydroxyl group interacts with the side chain of Asn2756. The remainder of the pantetheine tunnel is rather hydrophobic, with Phe2866, Leu2867, Phe2884, Phe2888, Leu2928, and Tyr2957 all bordering the channel through which the pantetheine approaches the active site. Of note, the first four residues are located on the lid loop, illustrating its contributions to the functional interaction with the PCP.

**Figure 4.**
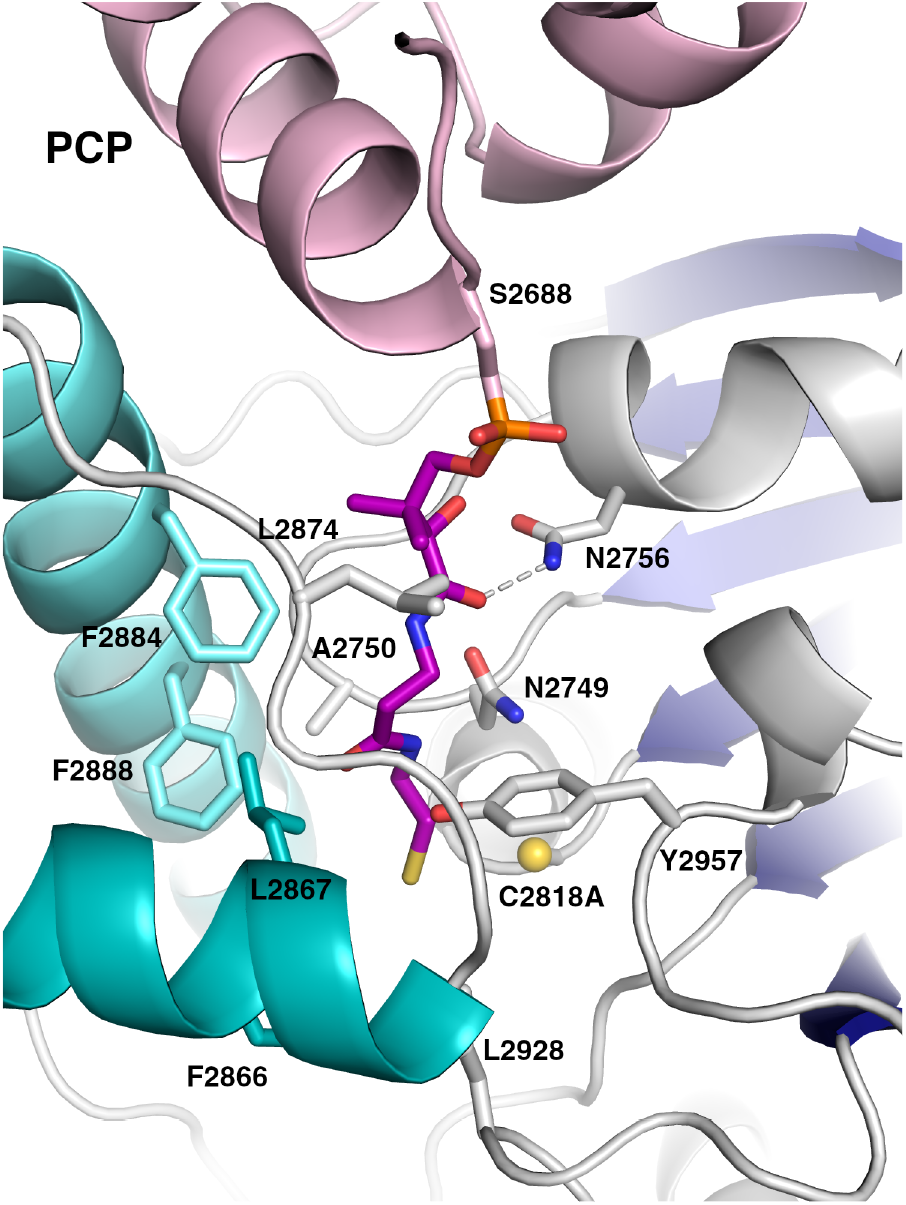
Pantetheine tunnel of the SulM thioesterase active site. The SulM thioesterase domain binds the PCP, allowing the pantetheine to pass through a mostly hydrophobic tunnel to enter the active site near the catalytic cysteine Cys2818, which has been mutated to an alanine for the complex structure.

### The interface between PCP and thioesterase domain

Acyl carrier protein domains generally contain four α-helices; three (α1, α2, and α4) are of similar length while α3 is shorter and lies perpendicular to the others (11,43). The pantetheinylation site is located at the N-terminus of helix α2. To enable the pantetheine to approach the active site, the PCP domain binds into a cleft on the surface of the thioesterase domain. In the SulM didomain protein, two regions of the PCP are responsible for interactions with the thioesterase domain. A hydrophobic patch on helix α3 containing Val2704, Thr2705, and Tyr2708 interacts with a complementary hydrophobic region on the thioesterase domain composed of Met2733 on strand β2 and Pro2754, Val2755, and Val2758 that lie on the helix between strands β3 and β4. A second series of interactions occurs with PCP helix α2 that follows the pantetheine site. Phe2689 in the α2 of PCP interacts with a hydrophobic pocket formed by Leu2874, Pro2875 and Phe2884 in the TE domain, as well as the dimethyl group of the pantetheine cofactor. The PCP also forms polar interactions through Lys2690 and Arg2693 of α2 with Asp2879 and Gln2877 of TE domain.

### Comparison of the PCP-TE complex with EntF

To date, only the EntF thioesterase domain has been structurally characterized, both by NMR (15) and crystallography (16), bound to its carrier protein partner in a functional interaction. Both structures show a similar core thioesterase fold and interact with the PCP. The PCP of the NMR structure contains a mutation of the pantetheinylation site serine to an alanine, while the crystal structure contains an analog of the acyl pantetheine and thus sits deeper in the thioesterase domain pocket. The PCP-thioesterase complex from EntF was captured crystallographically using an α-chloroacetyl amino phosphopantetheine designed to stabilize the inter-domain interaction (16). We superimposed the EntF and SulM thioesterase domains, allowing the comparison of the orientation of their respective PCP domain partners. The two thioesterase domains overlay with a rms displacement of 2.7 Å over all Cα positions excluding the flexible lid loops. The Cα positions of the serine residues from the two PCPs that harbor the pantetheine are 3.1 Å apart; however, the relative orientation of the two carrier domains differs. After superposition of the thioesterase core domains, the angle between the two α2 helices is 50° (Figure 5). Examining the size of the interface, the EntF interface is more substantial, covering 915 Å^2^ in area. In contrast, the interface between the PCP and thioesterase domain of SulM is 516 Å^2^.

**Figure 5:**
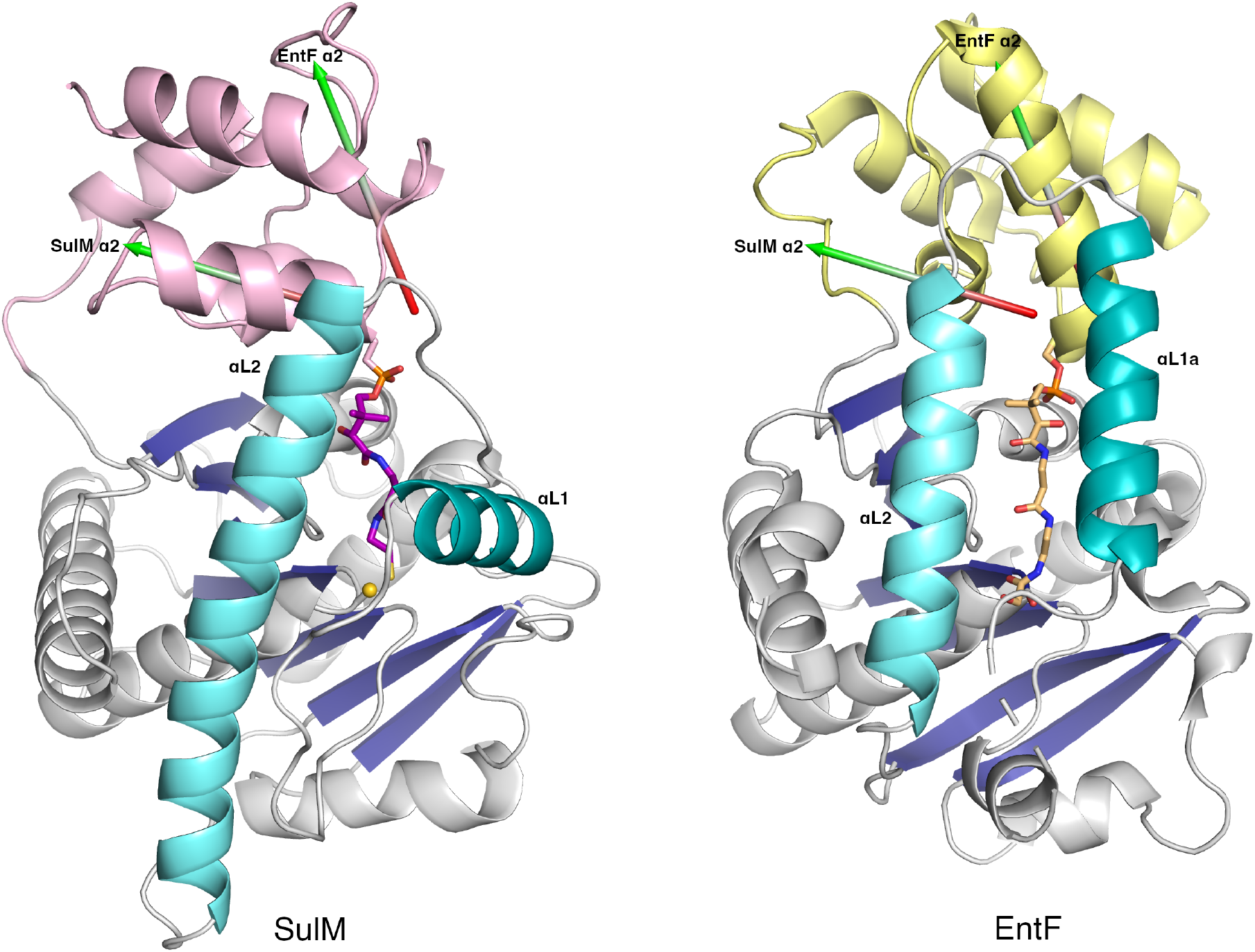
Comparison of SulM_PCP-TE and EntF_PCP-TE interfaces. The structures of the PCP-thioesterase didomain complexes were superimposed on the basis of the thioesterase domain, excluding the dynamic lid loops. The *holo*-PCP domains of each complex are represented in pink (SulM) or yellow (EntF). The PCP domains bind in the same region but adopt a different orientation. The α2 helix of each PCP is depicted with an arrow that is colored red to green in the N-to C-direction. The angle between the two helices is 50°, depicting the alternate positions adopted by the PCP domains.

Despite these differences, the panthetheine moieties adopt a similar orientation to enter the active site. The sulfhydryl of the SulTE pantetheine and the corresponding amine of the pantetheine analog in the EntF structure (16) are 2.5 Å from each other. The lid loop helices appear to be a major cause of the alternate positioning of the PCP domains. The αL2 helices are similar in positioning although, as noted previously, the helix of SulTE extends several turns longer at the C-terminal end. The positions of the preceding helices, αL1 of SulTE and αL1b of EntF, adopt very different orientations, with the αL1b helix of EntF running parallel to the αL2 helix. The C-terminal end of the αL1b helix and the loop that connects it to the αL2 helix would clash with the PCP position adopted in the SulTE structure, thereby pushing it away from the active site to adopt the orientation seen in EntF.

### The active site of β-lactam/lactone forming thioesterase domains enables proper orientation of the terminal nucleophilic residue

NRPS thioesterase domains carry out a variety of reactions to catalyze product release. In general, the reactions proceed through an initial transesterification reaction in which the peptide is transferred to the catalytic serine or cysteine. From here, the peptide can be released through a simple hydrolysis step or often through more complex macrocyclization reactions with nucleophilic groups from the peptide that may include side chains or the N-terminus of the peptide. Additionally, some thioesterase domains catalyze iterative peptide linkages in which two or more peptides produced by the upstream NRPS are combined before release of the linear or cyclic peptide multimer (6).

While macrocyclizations of NRPS products are relatively common, the SulM TE reaction poses the different challenge of forming the β-lactam ring, requiring that the enzyme has a constrained binding pocket. Biochemical evidence (23) shows that *N*-sulfonation by SulN precedes ring closure. Thus, to enforce 4-membered ring formation, that the enzyme must position the *N-sulfo-*β-amine of the peptide in a confined volume to allow nucleophilic attack (Figure 6A).

**Figure 6.**
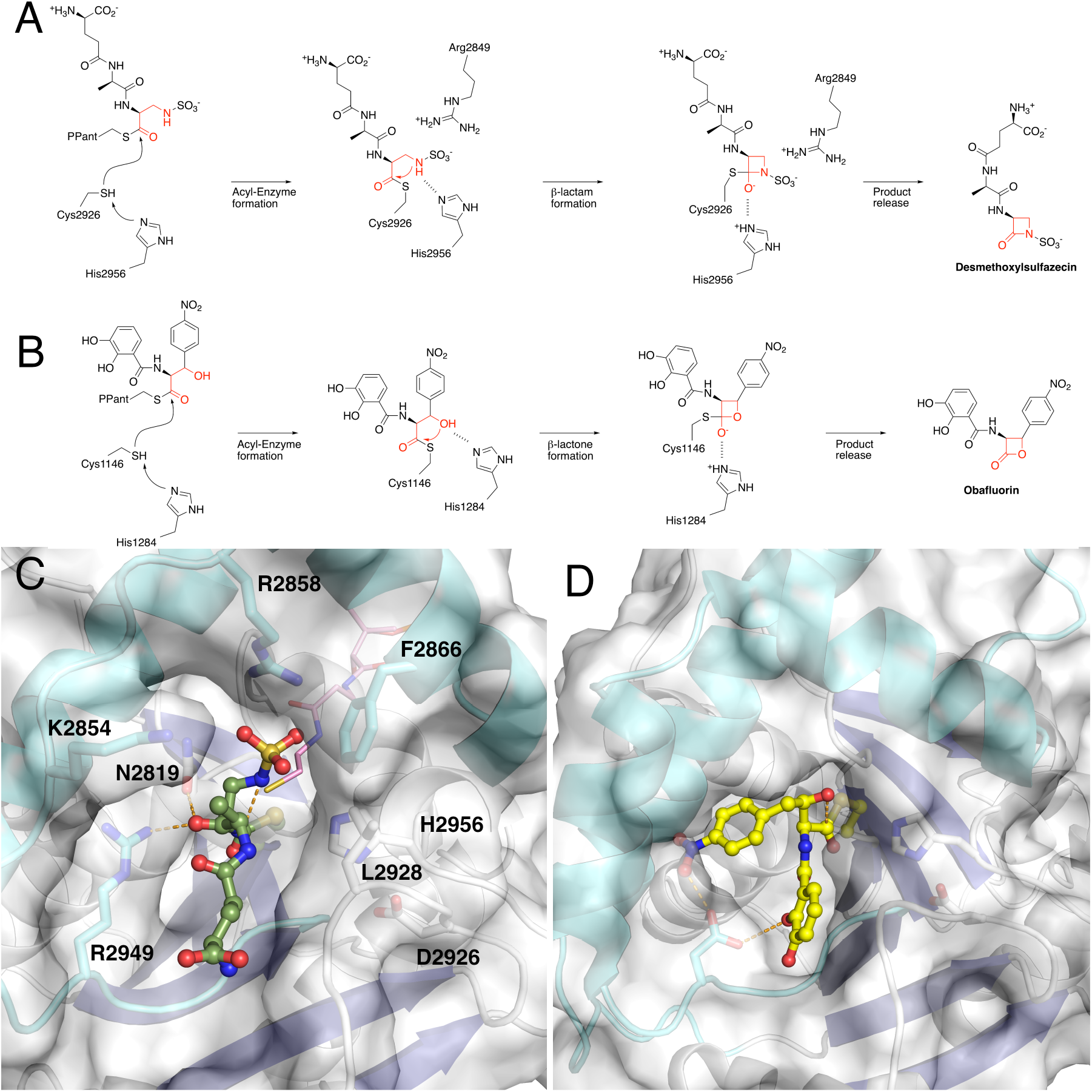
The thioesterase domain creates a cavity for β-lactam/β-lactone formation. Multistep reaction catalyzed by A) the SulM thioesterase and B) ObiF1 thioesterase domain. C. Active site of the thioesterase domain with a modeled linear sulfazecin peptide that cradles the substrate for β-lactam formation. Shown in pink is the pantetheine arm observed in the SulM PCP-TE structure. D. Active site of the thioesterase domain of ObiF1 (PDB **6N9E**) that positions the modeled obafluorin peptide for β-lactone formation. The view in panel D is rotated slightly around the Y-axis relative to panel C.

We modeled the sulfonated peptide into the active site with the goal of positioning the amine for attack and providing reasonable geometry for the remainder of the peptide given the relatively tight fit of the active site (Figure 6C). In this configuration, the side chain of Arg2858 is directed towards the sulfamate SO_3_, directing the nitrogen for attack on the thioester carbonyl. The nitrogen of the *N*-sulfoDAB residue is directed at the histidine residue of the catalytic triad, His2956. The proton on the sulfamate nitrogen has a pKa of ∼8 (44). Combined with the lack of additional residues in the active site capable of playing a role in deprotonation, this dual role of His2956 is the most plausible mechanistic model. The methylene groups of the Lys2854 side chain and Phe2866 further constrict the active site pocket. The carbonyl of the alanine residue from the sulfazecin tripeptide interacts with both Arg2948 and Asn2819, the residue that follows the catalytic cysteine at Cys2818 (Figure 6C).

Structure-based sequence alignment showed the Gln-Cys-Asn residues that border the catalytic cysteine are present only in SulTE compared to all other known thioesterase domain sequences from NRPS and PKS proteins (Figure S6). In contrast to the SulM thioesterase domain, all other domains have a hydrophobic residue in place of Asn2819. This Asn2819 residue following the catalytic Cys2818 is positioned near the C-terminal residue of the tripeptide in the docked tripeptide structure of PCP-TE. Thus, Asn2819 could interact with L-DAP residue in the native substrate indicating a unique role in β-lactam ring formation. Surprisingly, SulTE also showed a highly positively charged substrate binding pocket required to accommodate negatively charged glutamate and sulfamate groups of the tripeptide substrate (Figure S7).

We similarly modeled the active site of the ObiF1 thioesterase domain containing the peptide poised for formation of the β-lactone bond (Figure 6B, 6D). The pocket is similarly constrained to position the β-hydroxyl of the *p*-NO_2_-homophenylalanine moiety for attack on the carbonyl of the thioester. In the ObiF1 thioesterase domain model, Asp1177 contributes interactions to both the *p*-NO_2_ and hydroxyl of each aromatic ring. The remainder of the pocket is very hydrophobic in nature, simultaneously orienting the peptide while preventing access of water for hydrolysis of the thioester on the active site cysteine residue.

### Phylogenetic analysis and structure-guided genome mining

A phylogenetic tree of thioesterase domains selected from literature was generated using a Clustal Omega sequence alignment. The domain amino acid sequences were aligned with SulTE homologous sequences and a neighbor-joining phylogenic tree was constructed (Figure S8). Surprisingly, the β-lactam and β-lactone forming SulTE and ObiF1_TE grouped with Type-II proof-reading domains in the phylogenetic tree, which also have the catalytic Asp residue shifted to position II after β7 strand.

In order to identify other clusters harboring SulTE-like β-lactam forming TE domain, a PSI-BLAST search was performed using the SulTE sequence. Most of the hits with more than 70% similarity matched to SulM homologs from *Burkholdaria* bacteria with similar module organization previously suggested to synthesize sulfazecin (22). However, hits with lower than 70% sequence similarity identified clusters with different domain organizations in bacteria other than *Burkholdaria* and variable adenylation domain Stachelhaus code residues compared to the Sulfazecin cluster. To support the presence of β-lactam forming TE domains within these clusters, we established four criteria based on SulTE structural features (Figure 6 & Figure S9). The initial criterion involved examining for the presence of a catalytic triad residues with aspartate at position II and the presence of the Gln-Cys-Asn motif housing the catalytic cysteine, allowing for interaction of the Asn with the nucleophilic nitrogen. Additionally, TE domain sequences were distinguished by identifying positively charged residues at two out of three arginine positions, providing a positive active site environment to interact with the sulfamate. The presence of Stachelhaus residues that match the 2,3-diaminopropionate activation in the adenylation domain of the last module was considered an additional criterion as the DAP residue is essential to form the β-lactam ring. Finally, the presence of a sulfotransferase gene in the cluster was also considered.

We highlight three clusters that were chosen that varied from sulfazecin in the predicted amino acids activated by adenylation domains in upstream modules. On the basis of similar Stachelhaus code residues in the adenylation domain of the last module, these three clusters also most likely activate DAP for β-lactam ring formation. These clusters most likely biosynthesize peptides longer than sulfazecin indicating the synthesis of novel β-lactam products. However, further investigation is required to confirm β-lactam formation by these clusters. A cluster from *Flavobacterium* sp. Leaf82 bacteria (Figure 7) showed a large NRPS cluster predicted to be composed of seven modules. The upstream modules are predicted to produce a Orn-Gly-Asp-Asp-Xxx-Thr-DAP peptide. The fifth module showed the presence of an unknown domain in place of an expected adenylation domain, which matched a FkbH-like domain upon BLAST search and most likely incorporates an unknown monomer. A second biosynthetic gene cluster from *Chryseobacterium* sp. G0240 contained four modules including an epimerization domain in third module and is likely to place a D-ornithine preceding the β-lactam ring-forming DAP residue.

**Figure 7.**
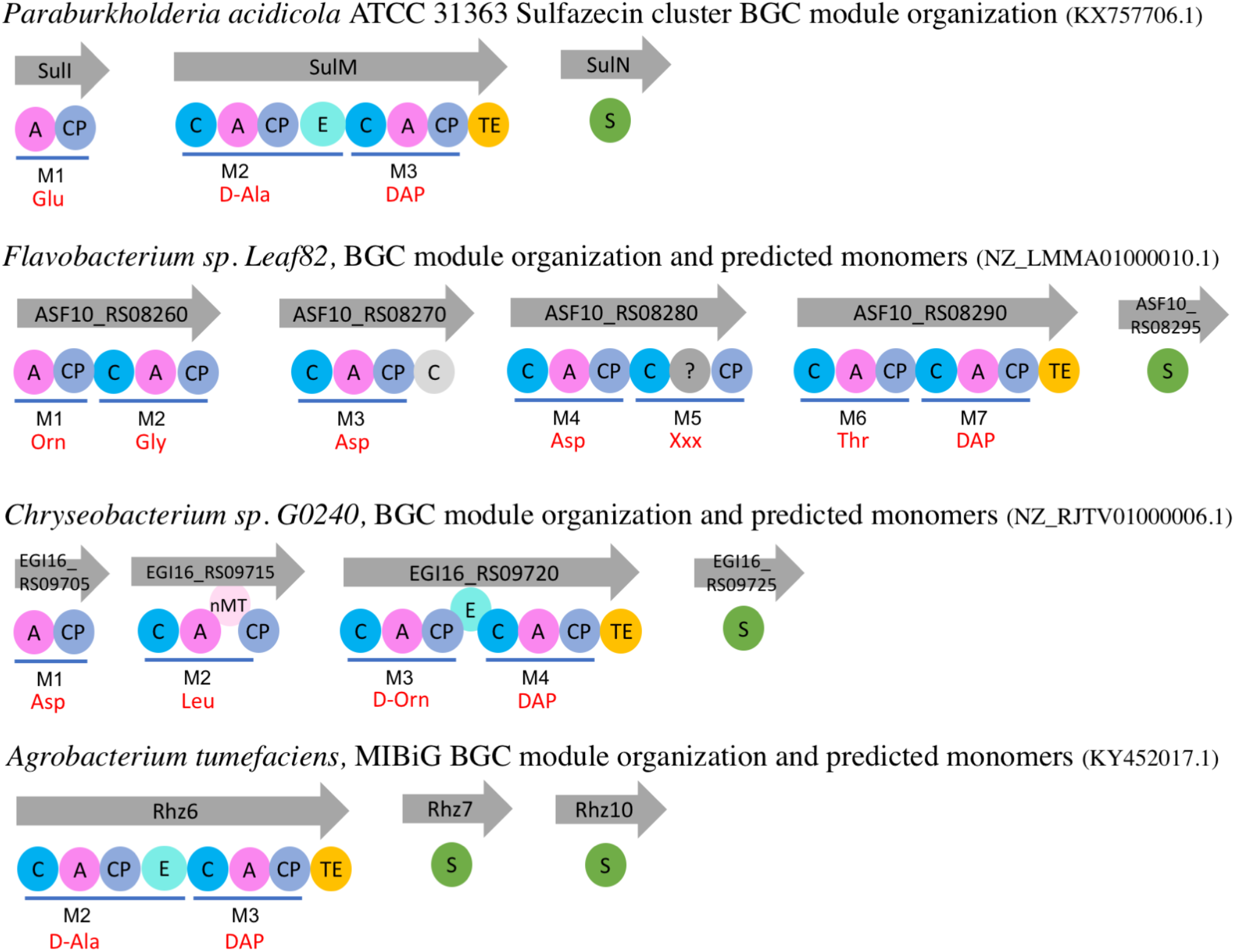
Anti-SMASH analysis of likely new beta-lactam forming clusters. Anti-SMASH 7.0 was used to search for biosynthetic clusters (BGCs) using SulTE sequence as query in PSI-BLAST. The BGCs harboring SulTE-like domains were analyzed for module architecture and predicted substrates for each module. The predicted adenylation domain substrates differed from Sulfazecin cluster except in last module which activates DAP required for β-lactam ring formation. Accession code for each cluster is provided in brackets and for each protein on the arrow. Predicted product and Stachelhaus code residues for adenylation domain in last module are shown in supplemental table S3. Figure legends are as followed, A: adenylation domain, CP: carrier protein, C: condensation domain, C in gray: probable condensation domain, E: epimerization domain, TE: thioesterase domain, nMT: methyltransferase domain, ?: FkbH-like domain, S: sulfotransferase domain.

Finally, examination of the MiBIG repository (1), identified an unpublished β-lactam-containing peptide from *Agrobacterium tumefaciens* that showed similarity to the SulTE domain sequence and satisfied the structure-guided criteria of the thioesterase domain and the presence of a sulfotransferase gene in the cluster (https://mibig.secondarymetabolites.org/repository/BGC0001671/index.html). This example also strongly supports the use of the SulM thioesterase structural features to identify new β-lactam harboring peptide products. Nevertheless, considering the differences in the sequence of the predicted peptide products, it is important to experimentally determine the identity and function of these peptides.

## Conclusions

We present here the structure of the β-lactam producing thioesterase domain from SulM on its own and as a complex with the upstream *holo*-PCP domain. The structure illustrates that despite the unusual chemistry catalyzed by the SulM thioesterase domain, the overall fold of the protein is broadly shared with other structurally characterized NRPS thioesterase domains. While the core thioesterase domain fold is adopted by SulM, the dynamic lid helices adopt an unusual conformation seen in the β-lactone forming thioesterase domain of the obafluorin biosynthetic protein, ObiF1 (39). We propose that the configuration of the lid helices results in a constrained active site that, upon formation of the acyl-enzyme intermediate, forces the *N*-sulfonated DAB moiety to adopt a geometry that is primed for β-lactam formation. This confined active site is also observed in the thioesterase domain of ObiF1.

The structure of the SulM thioesterase domain in complex with the upstream PCP is to our knowledge the second structure of the active complex between a PCP and the thioesterase domain. The complex is rotated ∼50° from the interface observed in the structure of EntF; however, both thioesterase domains direct the same face, the β2 strand, the helix that joins the β3 and β4 strands, and the lid helices towards the carrier protein and the trajectory of the pantetheine still allows a common approach to the active site.

Overall, these analyses indicate that apart from sulfazecin there might be more NRPS derived β-lactam antibiotics biosynthesized by bacteria and the importance of genome mining to discover novel natural products.

### Experimental Procedures

#### Cloning, Expression, and Protein Production

The accession code for SulTE and PCP-TE sequence in SulM is AOZ21320.1. The protein sequences of the expressed constructs are described in Figure S10.

The expression construct and purification strategy for PCP-TE, encoding Glu2660 through Ala2723, was previously described (23). The catalytic cysteine of the PCP-TE construct was mutated to an alanine. A final gel filtration run was used for structural analysis to remove aggregates and transfer the protein to storage buffer (20 mM MES, pH 6.0, 50 mM NaCl, and 0.5 mM triscarboxyethyl phosphine (TCEP)).

To produce an expression plasmid for the TE domain alone, multiple sites upstream of SulTE were screened for stable protein production. Oligonucleotides TE3-F and TE3-R were used to amplify the TE domain, which was initially cloned into pBluescript to give pBS/TE3.

TE3-F:5’-GGCATATGGCCTCGGAGGAGTCGAGCTCGATCGTG-3’

TE3-R 5’-GGAAGCTTTCTGCCGTCACACCTTTGCAGGACAC-3’

From pBS/TE3, the TE3 gene was excised with NdeI and HinDIII and ligated into pET28b. The final protein construct pET28B/TE3 encoded residues Ala2723 through Ala2984 for the TE domain.

Proteins were expressed in *E*. coli Rosetta2 (DE3). An overnight culture was used to inoculate a large-scale growth, which was grown to OD_600_ = 0.6-0.7 and cold-shocked on ice for 40 min before induction with 0.5 mM IPTG. The expression was carried out at 16 °C for 24 h. Cells were harvested by centrifugation. 10 g of cells were suspended in 50 mL lysis buffer (50 mM NaH_2_PO_4_, 10 mM imidazole, 300 mM NaCl, 10% glycerol, pH = 8.0). Lysozyme was added to the final concentration of 2 mg/mL and incubated on ice for 30 min. The treated cells were subjected to sonication at 40% AMP for 13 min with 9.0 sec/9.0 sec pulse (Model GEX 400 by Ultrasonic Processor). The cell debris was removed by centrifugation at 41400 × g for 25 min. The cell-free-extract was mixed with 3-5 mL Ni-NTA resin and incubated at 4 °C for 60 min. The resin was washed 2 × with 10-15 mL wash buffer (50 mM NaH_2_PO_4_, 20 mM imidazole, 300 mM NaCl, 10% glycerol, pH = 8.0). The TE3 was then eluted with 3 mL imidazole elution buffer (50 mM NaH_2_PO_4_, 250 mM imidazole, 300 mM NaCl, 10% glycerol, pH = 8.0). A final purification step on a gel filtration column into storage buffer (50 mM MES, pH 6.0, 50 mM NaCl, and 0.5 mM TCEP) was performed prior to crystallization.

#### Attachment of sulfazecin tripeptide mimic to PCP-TE

The endogenous sulfazecin tripeptide was mimicked using L-glutamate in place of L-DAP at the C-terminal position. The γ-D-Glu-D-Ala-L-Glu mimic was synthesized using conventional peptide chemistry to join D-Glu-D-Ala (23) to L-Glu. Peptidyl-mimic-S-CoA thioester was generated by coupling the tripeptide mimic to CoA (see supplementary methods). The CoA thioester was used to convert mutant *apo*-PCP3-TE C2818A to its *holo*-form by combining with the peptide CoA thioester with 2 µM of the promiscuous phosphopantetheinyl transferase Sfp (23). The loading reaction was incubated at room temperature for 1 h (21), then buffer exchanged into buffer by three serial dilutions and concentrated using an Amicon Ultra 3K (Millipore). Loading of the peptide mimic was confirmed by high resolution MS (Figure S11).

#### Crystallization and Structure Solution

Initial hits for crystallization of both proteins were identified from a high-throughput sparse matrix screen (Hauptman-Woodward Institute, Buffalo, NY) (45,46). Further optimization of crystallization was performed in micro-batch under oil plates at room temperature. A drop of 1 µL of protein was added by 1 µL of crystallization cocktail in each well and the plate was overlayed with 5 ml of mineral oil. Data collection for both protein crystals were performed remotely at SSRL beamline 9-2.

Crystallization of the free thioesterase domain was performed at 10-12 mg/ml with a crystallization cocktail containing 100 mM Bis-Tris Propane pH 7.0, 100 mM ammonium bromide, and 40% PEG 20,000. Initial crystals were long thin rods nucleating from a single point. These crystals were used as seeds to grow diffraction quality crystals in the same conditions. Cryo-protection of crystals was done by serial transfer into crystallization cocktail supplemented with 10 and 20% NDSB mixture (60% ethylene glycol and 200 mg/ml NDSB-201, in water). Diffraction data were indexed, scaled, and integrated resolution with HKL2000 in space group *P*2_1_. Structure determination was performed with Phaser in PHENIX using ObiF1 (39) thioesterase domain (PDB **6N8E**, residues 1059-1303) as molecular replacement model. The model was built and refined iteratively using COOT and PHENIX, employing Translation-Libration-Screw (TLS) parameters in final stages of refinement.

The didomain PCP-thioesterase construct was crystallized at 10 mg/ml with a cocktail containing 1.8M tribasic ammonium citrate, pH 7.0-7.5. Clusters of needle-like crystals were used as a seed stock to grow thin plate-like crystals. Further seed stock was made from these plate-like crystals and seeded into drops with different pHs from 7.0 to 7.5, made by mixing different ratios of stock solutions at pH 7 and 7.5. Crystals were transferred to fresh drops and diffraction quality crystals were obtained. Cryo-protection of crystals was done by transferring into mother liquor supplemented with 10% glycerol. From a single crystal that grew from a cocktail containing a 70:30 cocktail ratio of pH 7.0 and pH 7.5, diffraction data were collected, then indexed, scaled, and integrated with HKL2000 in space group *P*2_1_. The single domain SulM thioesterase structure was used as molecular replacement model for structure determination with Phaser in PHENIX. Model building and refinement was done iteratively using COOT and PHENIX. The PCP domain was built manually in COOT. The phosphopantetheine was modeled in the observed density and refined. Although loaded, the density for the peptide mimic was not of sufficient quality to be included in the model Translation-Libration-Screw (TLS) parameters were used during final stages of refinement.

##### Computational

The superposition of SulM thioesterase domain with existing NRPS structures was performed with PYMOL using the super command and the full domain lacking the lid loop spanning β6 and β7. The root mean square (rms) displacement is reported in the supporting information. The topology diagram was adapted from Horsman *et al*., (6) and was created with TOPDRAW (47). The angle between helices module of PYMOL is provided under open access license at: https://pymolwiki.org/index.php/AngleBetweenHelices.

The phylogenetic tree generation was calculated using structurally characterized thioesterase domain sequences, which were aligned in Clustal Omega. The phylogenetic tree was downloaded and visualized in iTOL (48). A bioinformatic search for potential *β-lactam forming clusters* was performed by using the SulM protein sequence in PSI-BLAST search for homologous sequences. Hits lower than 70% similarity were analyzed for presence of other NRPS modules in the cluster, and the identification of the structure-guided sequence motifs and a sulfotransferase gene, as described in Results.

## Data Availability

The atomic coordinates and structure factors used to solve the structures of SulM TE (**8W2C**) and the didomain SulM PCP-TE (**8W2D**) have been deposited with the wwPDB.

### Acknowledgement

The investigation was supported by NIH grants GM136235 (to AMG) and NIH AI014937 and AI121072 (to CAT). Use of the Stanford Synchrotron Radiation Lightsource, SLAC National Accelerator Laboratory, is supported by the U.S. Department of Energy, Office of Science, Office of Basic Energy Sciences under Contract No. DE-AC02-76SF00515. The SSRL Structural Molecular Biology Program is supported by the DOE Office of Biological and Environmental Research, and by the National Institutes of Health, National Institute of General Medical Sciences (P30GM133894). Crystallization screening at the National Crystallization Center at HWI was supported through NIH grant R24GM141256.

## Supporting Information

Supporting information is available for this article.

## Author Contributions

The experimental crystallography described in the manuscript was performed by K.D.P. Synthetic chemistry was carried out by R.A.O. and M.S.L. Protein production was conducted by R. F. L, R.A.O, and K.D.P. Data analysis and interpretation, and manuscript preparation were done by K.D.P., R.A.O., R.F.L., C.A.T., and A.M.G.

