## Supporting Information for "The structure of the monobactam-producing thioesterase domain of SulM forms a unique complex with the upstream carrier protein domain"

### General Synthetic Methods

All chemicals and reagents were purchased from Sigma Aldrich (St. Louis, MO), AK Scientific (Union City, CA), or Fischer Scientific (Hampton, NH), unless otherwise indicated, and used without further purification. All solvents were distilled before use, including THF (Na / benzophenone), DCM (CaH), and MeCN (4Å MS). Anhydrous DMF was purchased from Sigma Aldrich and used without further purification.

Preparative HPLC methods were carried out on an Agilent 1100 (Santa Clara, CA) series HPLC equipped with a multi-wavelength UV-Vis detector using one of the following HPLC prep methods:
HPLC Preparatory Method A: [binary gradient: water +0.1% TFA (solvent A), acetonitrile +0.1% TFA (solvent B), 4.5 mL/min]: 0-30 min gradient 0% to 100% B; 30-31 min gradient 90% to 10% B; 31-35 min isocratic 10% B. Phenomenex Kinetex® 5 µm C18 100Å LC column 250 10 mm.
HPLC Preparatory Method B: [binary gradient: water +0.1% TFA (solvent A), acetonitrile +0.1% TFA (solvent B), 4.5 mL/min]: 0-30 min gradient 0% to 60% B; 30-31 min gradient 90% to 10% B; 31-35 min isocratic 10% B. Phenomenex Kinetex® 5 µm C18 100Å LC column 250 10 mm.

UPLC-HRMS experiments to determine exact masses and purity levels of organic compounds were carried out on a Waters Acquity / Xevo-G2 UPLC-MS system at the Johns Hopkins Mass Spectrometry Facility. NMR spectra were recorded on either 400 MHz or 300 MHz Bruker (Billerica, MA) Advance NMR spectrometers. Chemical shifts are reported relative to the reference shift for the solvent used relative to TMS. Many 1H-NMR resonances are broad owing to amide and carbamate configurational equilibria. 13C NMR spectra were recorded on a 400 MHz Bruker Advance operating at 101 MHz. Chemical shifts are reported relative to the reference chemical shift of the NMR solvent. In 13C experiments for which D2O is the solvent, an internal standard of acetone was added, and the spectrum adjusted to this reference (215.94 ppm, 30.89 ppm).

Reagent abbreviations are as follows:

PyBOP: (Benzotriazol-1-yloxy)tripyrrolidinophosphonium hexafluorophosphate
DCM: dichlorormethane
DMF: *N,N*-dimethylformamide
DIPEA: *N,N*-diisopropylethylamine
MeCN: acetonitrile
TFA: trifluoroacetic acid

Thin layer chromatography (TLC) was carried out on silica gel coated glass plates with the elution conditions indicated, CV refers to column volumes of mobile phase used in silica gel chromatography, and tR indicates retention time.

**Synthesis of γ-D-Glu–D-Ala–L-Glu-CoA Tripeptide mimic**

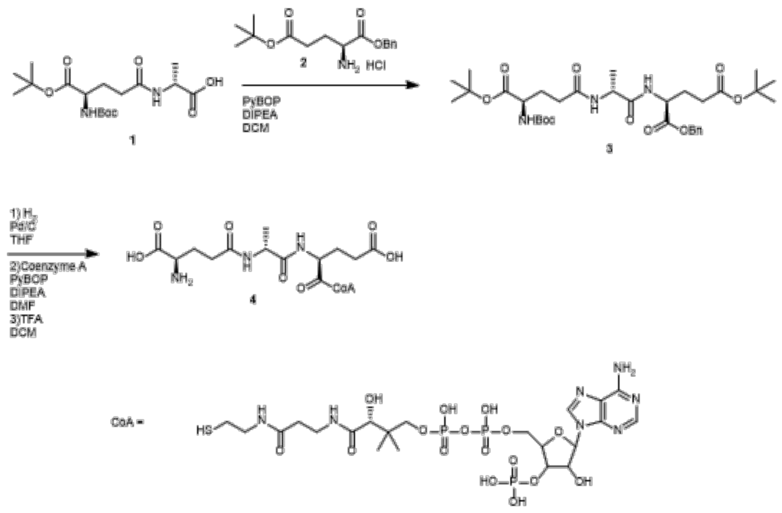

**Protected γ-D-Glu–D-Ala–L-Glu tripeptide benzyl ester 3**. Protected D-Glu-D-Ala dipeptide 1 (0.45 mol) was dissolved in 5 mL of DCM. To the solution was added doubly protected D-glutamate **2** (1.35 mmol) followed by PyBOP (0.54 mmol) and DIPEA (1.39 mmol) and the solution stirred overnight at room temperature. The organic phase was washed with aqueous $NaHCO_3$ and $NH_4Cl$, and the combined organics were dried over anhydrous $MgSO_4$. Protected γ-D-Glu-D-Ala–L-Glu tripeptide benzyl ester **3** (0.23 mmol 51 % yield) was isolated by silica gel column chromatography eluting with EtOAc/hexanes. **$^1$H-NMR** (400 MHz, $CDCl_3$): δ 7.38-7.28 (m, 5H), 7.03 (d, *J* = 7.8 Hz, 0.5H)6.49 (d, *J* = 6.5 Hz, 0.5H), 5.23 (d, *J* = 7.8 Hz, 0.5H), 5.18-5.08 (m, 2H), 4.61-4.53 (m, 0.5H), 4.54-4.45 (m, 0.5H), 4.16-4.08 (m, 0.5H), 3.49 (dd, *J* = 8.3, 5.2 Hz, 0.4H), 2.33 (t, *J* = 7.5 Hz, 1H), 2.29-2.20 (m, 2H), 2.20-1.74 (m, 4H), 1.56 (s, 11H), 1.44 (s, 5H), 1.42-1.39 (m, 13.5H), 1.36 (d, *J* = 7.0 Hz, 1.5H) **HRMS** (ESI) *m/z*: [M+H]+ $C_{33}H_{52}N_3O_{10}$ calculated 650.3647, found 650.3466

**γ-D-Glu–D-Ala–L-Glu-CoA Tripeptide mimic 4.** Protected γ-D-Glu-D-Ala-L-Glu tripeptide benzyl ester **3** (0.088 mmol) was dissolved in 10 mL anhydrous THF. Pd/C (5.7 mg, 10% wt/wt) was added and the solution stirred under balloon pressure hydrogen gas for 3 h at room temperature. The solution was then filtered through a 0.5 µM filter and the volatiles removed *in vacuo*. The resulting free acid peptide (0.063 mmol) was dissolved in 2.5 mL of anhydrous DMF. Coenzyme A sodium salt hydrate (0.069 mmol) and DIPEA (0.14 mmol) were added followed by PyBOP (0.075 mmol). The reaction was stirred under inert atmosphere of argon for 2 h at room temperature and then HPLC purified (HPLC method A) to yield protected γ-D-Glu–D-Ala–L-Glu-CoA tripeptide thioester. The lyophilized material was subsequently deprotected by stirring in 2 mL of a 3:1 TFA:DCM solution at room temperature for 1 h. The volatiles were then removed *in vacuo* and the reaction resuspended in 3 mL of ACN and purified by HPLC method B to yield γ-D-Glu–D-Ala–L-Glu-CoA tripeptide mimic **4** (0.0098 mmol, 11%). **$^1$H-NMR** (400 MHz, $D_2O$): δ 8.88 (s, 0.75H), 8.85 (s, 0.25H), 8.68 (s, 1H), 6.55 (d, *J* = 5.5 Hz, 0.25H), 6.44 (d, *J* = 5.5 Hz, 0.75H), 5.43-5.35 (m, 0.35H), 5.19-5.10 (m, 1.65H), 4.93-4.88 (m, 0.4H), 4.69-4.63 (m, 0.35H), 4.61-4.45 (m, 3H), 4.34-.428 (m, 1H), 4.25 (s, 1H), 4.15-4.08 (m, 1H), 3.92-3.84 (m, 1H), 3.72-3.66 (m, 2H), 3.61-3.55 (m, 2H), 3.29-3.22 (m, 2H), 2.82-2.74 (m, 2H), 2.74-2.64 (m, 4H), 2.52-2.39 (m, 3H), 2.24-2.12 (m, 1H), 1.68-1.61 (m, 3H), 1.18 (s, 3H), 1.07 (s, 3H). **HRMS** (ESI) *m/z*: [M+H]+ $C_{34}H_{56}N_{10}O_{23}P_3S$ calculated 1097.2448, found 1097.2446

Spectral characterization of synthesized compounds 3 and 4 are presented in Figures S11-13.

**Supplemental figures**

| Protein | PDB | SulM | SrfA-C | Vlm2 | EntF | NocB | ObiF | FenB | SkyXY | AB3403 |
|---|---|---|---|---|---|---|---|---|---|---|
| **SulM** | **8W2C** | | 21 % | 20 % | 22 % | 21 % | 25 % | 19 % | 19 % | 17 % |
| **SrfA-C** | **2VSQ** | 1.8 Å | | 22 % | 22 % | 23 % | 19 % | 34 % | 21 % | 22 % |
| **Vlm2** | **6ECE** | 3.2 Å | 1.4 Å | | 20 % | 23 % | 23 % | 20 % | 25 % | 26 % |
| **EntF** | **3TEJ** | 2.7 Å | 2.0 Å | 2.0 Å | | 22 % | 24 % | 26 % | 28 % | 25 % |
| **NocB** | **6OJD** | 2.4 Å | 1.4 Å | 1.5 Å | 1.6 Å | | 25 % | 24 % | 27 % | 27 % |
| **ObiF** | **6N8E** | 1.3 Å | 1.5 Å | 1.7 Å | 2.6 Å | 2.0 Å | | 21 % | 23 % | 21 % |
| **FenB** | **2CB9** | 1.8 Å | 1.1 Å | 1.4 Å | 1.9 Å | 1.3 Å | 1.6 Å | | 23 % | 21 % |
| **SkyXY** | **7CRN** | 2.9 Å | 2.0 Å | 1.7 Å | 1.2 Å | 1.2 Å | 2.0 Å | 1.6 Å | | 25 % |
| **AB3403** | **4ZXI** | 2.4 Å | 1.5 Å | 1.4 Å | 1.5 Å | 1.2 Å | 1.9 Å | 1.5 Å | 1.4 Å | |

**Supplemental figure S1. Sequence and Structure Alignment of SulM thioesterase domain compared with prior NRPS thioesterase domain structures**. Values above the diagonal represent pairwise sequence identity of the thioesterase domains, calculated with CLUSTAL OMEGA. Values below the diagonal represent rms displacement by superimposing the core thioesterase domain with PYMOL super algorithm, lacking the dynamic lid loops

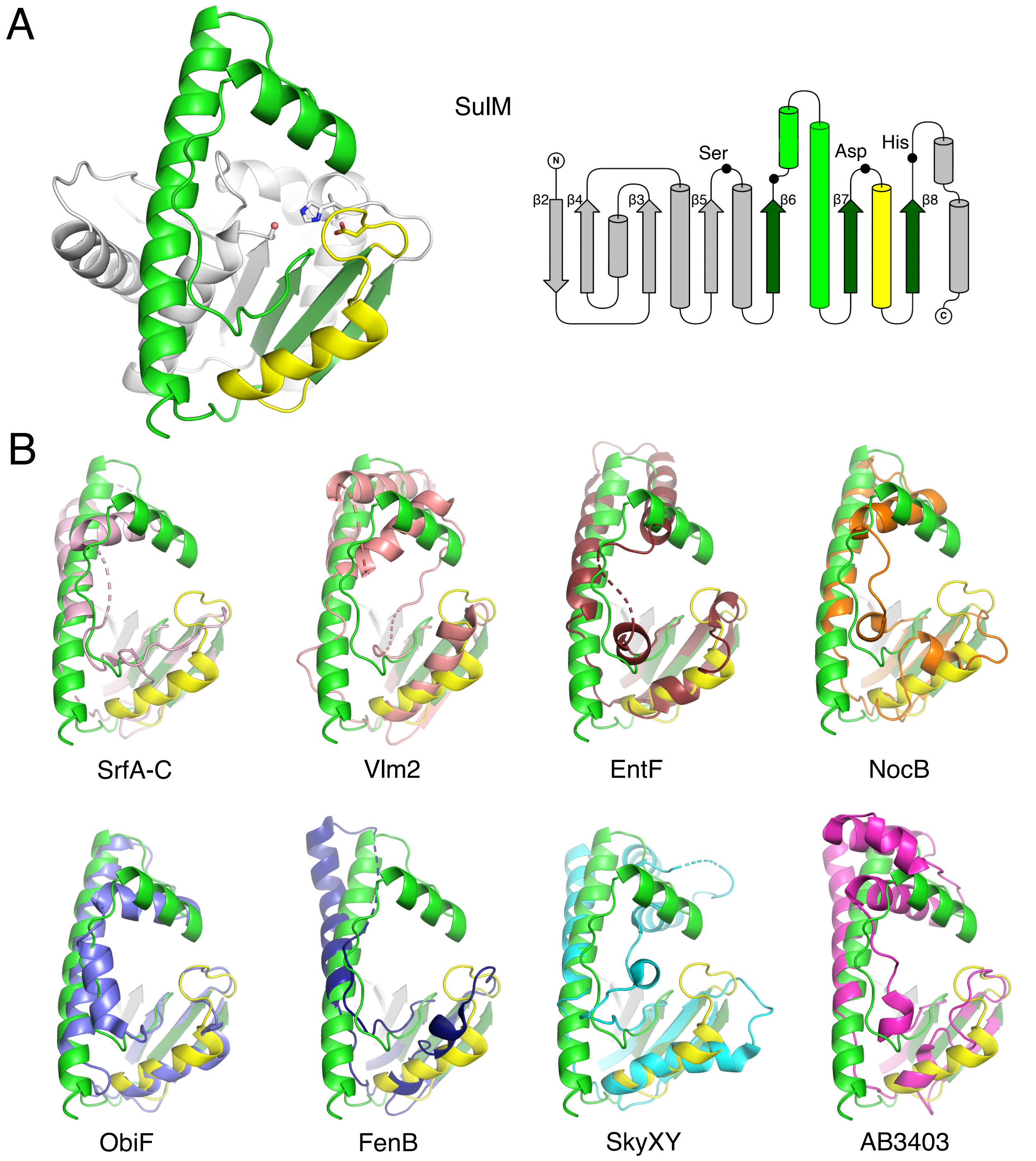

**Supplemental figure S2. Comparison of Lid regions between different NRPS thioesterase domain structures**. A. Ribbon diagram of SulM thioesterase. The side chains of the residues that form the catalytic triad are shown. The pink sphere reflects the use of the C2818A mutant enzyme. The green sphere in the ribbon and the unlabeled black circle in the topology diagram highlight the more common position of the Asp in NRPS thioesterase domains. Topology diagram created with TopDraw and adapted from M.E. Horsman et al., (2016). B. NRPS thioesterase domains (unique colors) were superimposed on SulM via the core of the protein. The overlay depicts the strands β6-β8 and the lid loop (light green) and the loop following β7 (yellow).

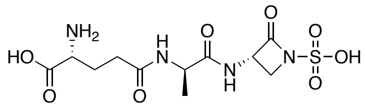

**desmethoxylSulfazecin**
**mw 366.4**

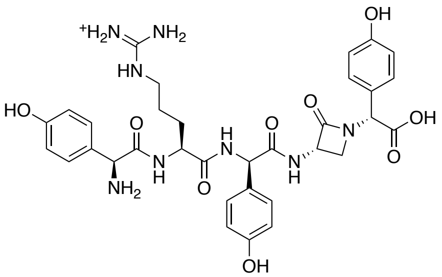

***p*-HPG-Arg-Nocardicin G**
**mw 691.7**

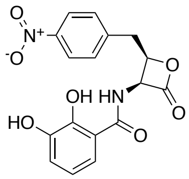

**Obafluorin**
**mw 358.3**

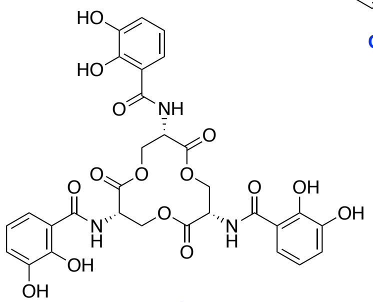

**Enterobactin**
**mw 669.5**

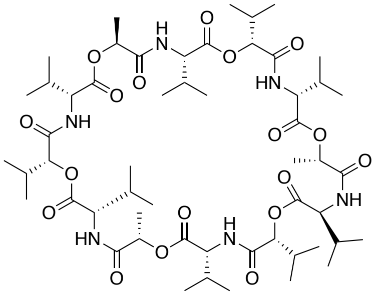

**Valinomycin**
**mw 1111.3**

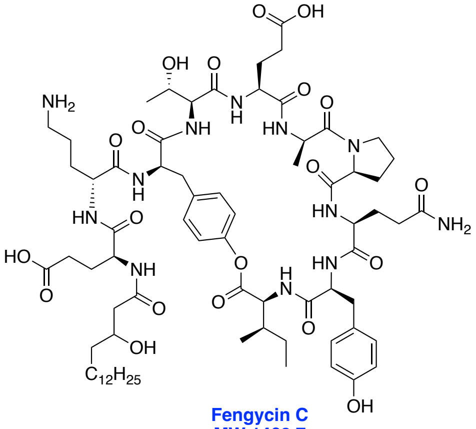

**Fengycin C**
**MW 1463.7**

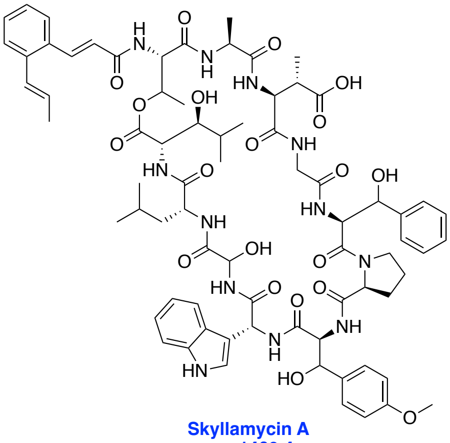

**Skyllamycin A**
**mw 1469.4**

**Supplemental figure S3. Chemical structures of final products released from thioesterase domains of structurally characterized proteins**. Release products of NRPS systems for which TE domain structure has been solved. In some cases, the product is further modified following release to yield the ultimate natural product. The structures and molecular weight do not include any chemical modifications that occur following release of the NRPS peptide.

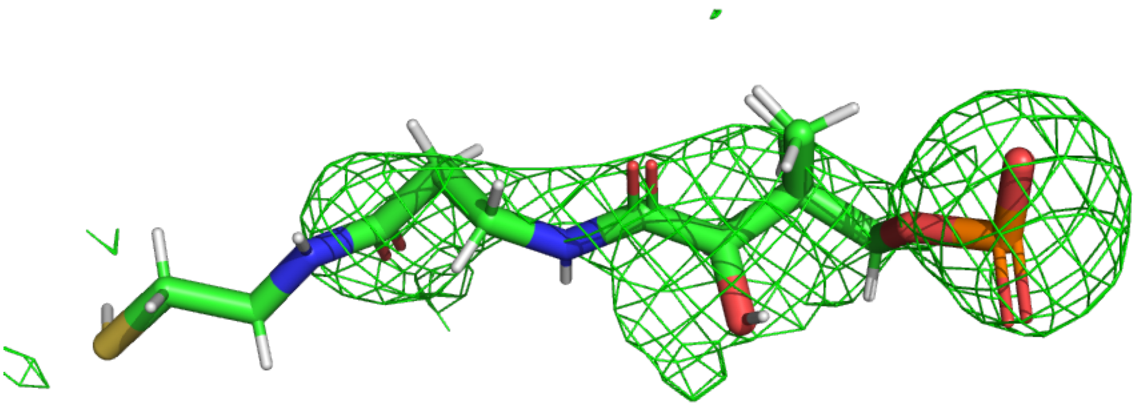

**Supplemental figure S4. The SulM PCP-thioesterase didomain structure illustrates the phosphopantetheine cofactor**. Omit map electron density of the phophopantetheine cofactor was produced by removing the phosphopantetheine group and submitting the final structure to a single round of simulated annealing refinement. The map is calculated with coefficients of the form Fo-Fc and contoured at 2.5σ.

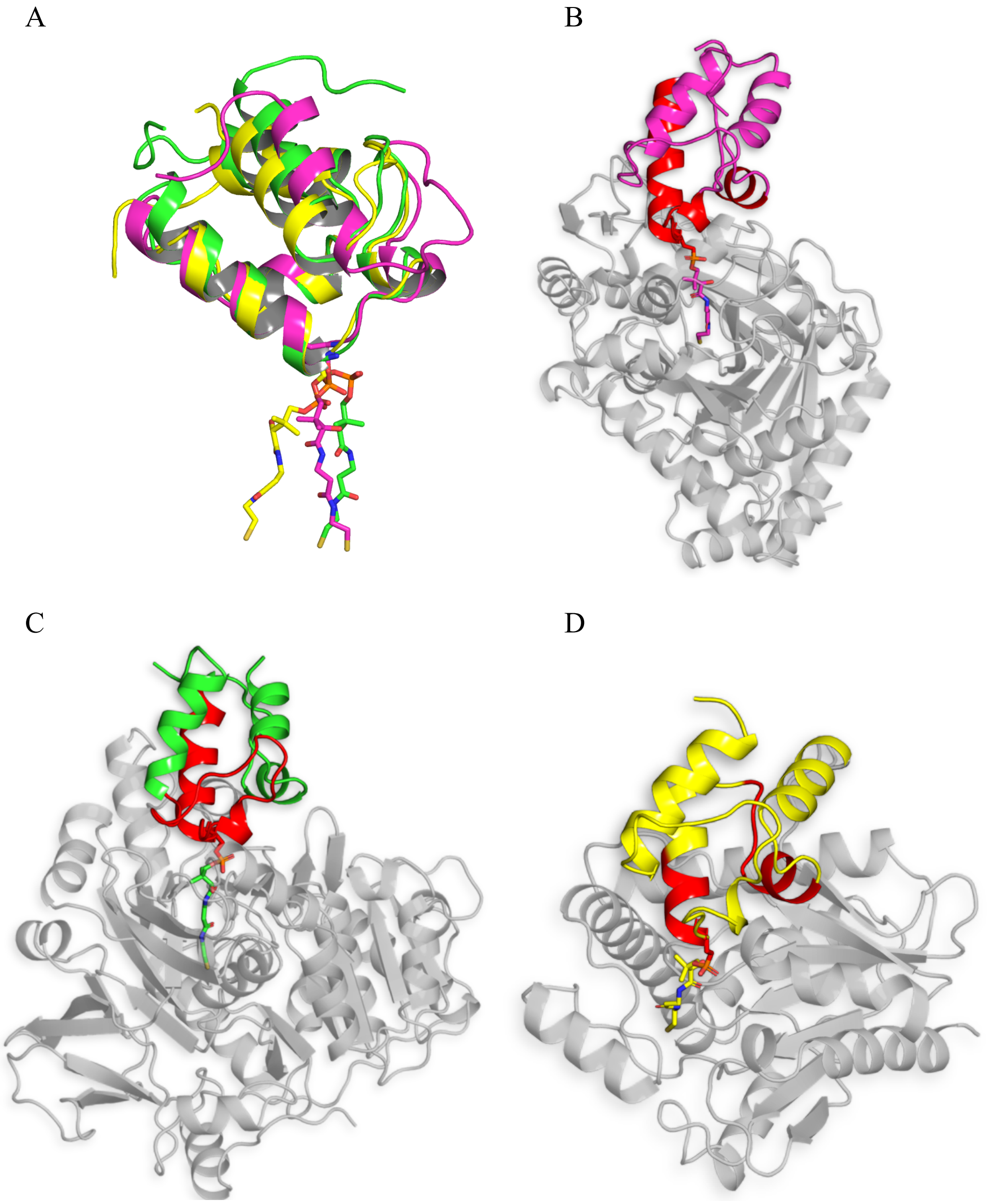

**Supplemental figure 5: Phosphopantetheine arm orientations and interaction interfaces of PCP with NRPS domains.** A, orientations of phosphopantetheine arm to approach the neighboring active sites in adenylation domain (pink, PDB **3RG2**), condensation domain (green, PDB **4ZXI**), and SulM thioesterase (yellow, PDB **8W2C**). Interfaces of carrier domain (red) while interacting with B. adenylation domain (pink, PDB **3RG2**), C. condensation domain (green, PDB **4ZXI**), and D. SulM thioesterase (yellow, PDB **8W2C**), highlighting the orientation of the cofactor adopts to approach the neighboring active site. The PCP interactions with three catalytic domains employ different regions of the carrier domain.

```
FengTE_2CB9__              42  ALQLN---HKAAVYGFHFI---------------EEDSRIEQYVSRITEIQ-PEGPYVLLGYSAGGNLAF   92
2cb9_chainA_p007           36  ALQLN---HKAAVYGFHFI---------------EEDSRIEQYVSRITEIQ-PEGPYVLLGYSAGGNLAF   86
7VJT_                      46  LGRLP---ADTPMYGFERVE-------------GSIEERAQQYVPKLIEMQ-GDGPYVLVGWSLGGVLAY   98
7vjt_chainA_p006           39  LGRLP---ADTPMYGFERVE-------------GSIEERAQQYVPKLIEMQ-GDGPYVLVGWSLGGVLAY   91
6BA8                       38  QAEQL---ADCALSLVTWPGRDRLRHL---EPLRSITQLAALLANELEASVSPDTPLLLAGHSMGAQVAF  101
TycC_TE                    20  AAEIQ----GVSLYSFDFI---------------QDDNRMEQYIAAITAID-PSGPYTLMGYSSGGNLAF   69
SrfAC_TE                   20  SSRLP----SYKLCAFDFI---------------EEEDRLDRYADLIQKLQ-PEGPLTLFGYSAGCSLAF   69
RifR_                      40  AKALA---PAVEVLAVQYPGRQDRRHE---PPVDSIGGLTNRLLEVLRPFG--DRPLALFGHAMGAIIGY  101
3flb_chainA_p003           36  AKALA---PAVEVLAVQYPGRQDRRHE---PPVDSIGGLTNRLLEVLRPFG--DRPLALFGHAMGAIIGY   97
6VAP_                      44  AAALA---PRCDVLAVQYPGRQDRRAE---KPLEDIDELANQLFPVLRARV--HQPVALFGHSMGATLAF  105
6FVJ_                      48  SREFS---ADVKRIAVQYPGQHDRSGL---PPLESIPTLADEIFAMMKPSARIDDPVAFFGHSMGGMLAF  111
7E3Z_                      43  AEDLP---EDVELALICYPGREARFGA---PFARVWTELRDDVVRSVRGLT--GRPYILFGHSMGSWMAF  104
3QIT_                      46  ALPLA--AQGYRVVAPDLFGHGRSSHLE-MVTSYSSLTFLAQIDRVIQELP--DQPLLLVGHSMGAMLAT  110
TrdC                       38  ASGLG---DLLPCSIWETIPQEAG-------TGAGDADPVERWLEEVAADG--RPVRAVFGFCVGGVYAG   95
LipX2                      47  GRLLA---RGVPLWETRQPEPEQA-------RTFGGEDFASYWVRGVRDTG--RPVRAVLGYCVGGLYAA  104
KSE_70420                  39  LPHLN---PDFSLWQTVAPVRAPE-------TGAPPEEYLAAWLSEIEASG--RVVRAVMGYCAGSVFAG   96
KirH1                      39  TPSLG---LDCALWQTVAPARAPG-------PGIDPDEYLAPWLAEVAASG--RRVRAVLGYCAGSVFAG   96
SlgL                       53  VPNLP---ADRTVWETTQPALGQE-------TEMGGAAYVARWMRAVTESG--RYVRAVMGYCVGSVFAA  110
3QMV_                      61  QERLG---DEVAVVPVQLPGRGLRLRE---RPYDTXEPLAEAVADALEEHR-LTHDYALFGHSXGALLAY  123
7R0X_                      70  AENLG---DDYAIWGIHSRGFLTRH-----PPLKSIESXANYYIEILQAGN-ATGPYELAGYSXGGIIAY  130
NocTE                      39  APLLP----DVRLHVLSDPRFGQSD-----NRFATLAEMATRYVEWVRTTE-PEGPYRLGGWSFGGVVAL   98
6ojd_chainA_p005           32  APLLP----DVRLHVLSDPRFGQSD-----NRFATLAEMATRYVEWVRTTE-PEGPYRLGGWSFGGVVAL   91
3LCR__                     81  AEELD---AGRRVSALVPPGFHGGQ-----ALPATLTVLVRSLADVVQAEV-ADGEFALAGHSSGGVVAY  141
FtdB_TE_                   89  AGALR---GIAPVRAVPQPGYEEGE-----PLPSSMAAVAAVQADAVIRTQ-GDKPFVVAGHSAGALMAY  149
Erya3_TE                   21  AGALR---GIAPVRAVPQPGYEEGE-----PLPSSMAAVAAVQADAVIRTQ-GDKPFVVAGHSAGALMAY   81
MlaG_TE__                  22  GSEFR---DVRPVSALALPGFQRGE-----PLPESVEVLSQVLGEAVLAAA-DGDPYVLLGYSSGGIIGH   82
HitP3_TE_                  22  AAHFR---GVRPVSALPLVGFARGD-----LLPATAAAAQVVAENVLRAA-GGEPFVLVGYSSGGVLAY   82
VinP4_TE                   22  AAHFR---GVREVSALPMPGFATGD-----LLPATAEAVVEVFGRSLLRAA-GDEPFVLLGYSAGGVFAH   82
PikTE                      91  STSFQ---EERDFLAVPLPGYGTGTGTGTALLPADLDTALDAQARAILRAA-GDAPVVLLGHSGGALLAH  156
TaaE_TE2                   20  TDAFG---PTWPIHGLQPRGLDAKA-----VPHSRVESAAQAYLVALEQQC-PSGPVHLLGHSFGGWVAF   80
MassC_TE2                  20  SEALG---RDWPLYGLQPRGLDGEA-----VPHSQVEAAAQCYLAALEQEC-PEGPVHLVGHSFGGWVAL   80
LybB_TE2                   20  TQACG---AHTPIIGLQARGHDGRS-----VPHASVEAAVEALLPEVRALG-QGGPYHLLGHSFGGWIAF   80
6ECB_                      39  ARHLK---GFGEIHGVEAPGLGAGET----PVYPSFEEMVQFCSDSAAGVA--GDGVYIGGHSLGGHIAF   99
BdObiF_TE                  34  ARALP---DGFACSALQLPGHDPAAPD---EAFVDLDTTIDRAVDRLLAEA--AAPIVVYGHCAGNALAV   95
BdObiF_TE.pdb_chainA_s001  34  ARALP---DGFACSALQLPGHDPAAPD---EAFVDLDTTIDRAVDRLLAEA--AAPIVVYGHCAGNALAV   95
LnmJ_TE_                    5  FAALS---ERYRVIVVHHPGVGDTTA----CEELGYEGIADLCLRALRRLG-VQGPVHVAGASFGGITAQ   66
3ILS_                      41  PR-LK---SDTAVVGLNCPYARDPE-----NMNCTHGAMIESFCNEIRRRQ-PRGPYHLGGWSSGGAFAY  100
3ils_chainA_p004           41  PR-LK---SDTAVVGLNCPYARDPE-----NMNCTHGAMIESFCNEIRRRQ-PRGPYHLGGWSSGGAFAY  100
SulM_TE                    45  SRAMPEQASDVAMFGVKLPRTEVDSDG---AMLEEVRRLSNAVCDDLLAAT--DLPAIVFAQCNGSALAL  109
SSHG_TE                    20  AAQLP----EYRVVAFNYLP---------------GDDKVARYADLVEAAR-PEGACRLLGYSLGGNLAF   69
FtdB                      138  ATSLP----SHRLIAFNYLP---------------GDDKVSRYADLVAATV-PEGPVPLLGYSLGGNLAF  187
SGR814_TE                  20  AAQLP----EYAVIGFNYLP---------------GDDKVARYADLIEAAR-PEGACLLLGYSLGGNLAF   69
Hsaf                       20  AVHLP----EYEFVSFNYLL---------------GDDKVARYADLIEGIH-AEGHCTLFGYSLGGNLAF   69
IkaA                       20  AARLP----EFEFLAFNYLM---------------GEDKVSGYADLVAGHR-PEGEIDLLGYSLGGNLAF   69
TaaE_TE1                   20  GQHLD---ADIPVYGLEGVAWGE-------PQLQTMECLAARHIDIMRGVQ-PHGPYRLAGWSFGGVLAY   78
MassC_TE1                  20  GQHLP---GDYPIYGLPGVALGE-------AHLDSMEGLAARMVGLIRQIQ-PHGPYRLAGWSFGGVLAY   78
LybB_TE1                   20  SRHFD---DDLPIYGIQADRADA-------QREVSAQSLARRYLEVIRSVQ-PQGPYRLAGWSGGGHLAY   78
HbnA                       20  AVHSR---SAEPLAAVSYPGEED-------GRPGTLRELAALYVEQIRREQ-PHGPYRLGGYSFGGNVLF   78
2K2Q_                      33  HAFLQ---GECEMLAAEPPGHGTNQT----SAIEDLEELTDLYKQELNLRP--DRPFVLFGHSMGGMITF   93
7CRN_                      43  MQHIG---ADRPLYGLQARGLADPSA----TLPSSIEEMAADYVTQIRGVQ-PSGPYHLLGWSLGSLVIH  104
Pys-Pflu_TE                20  LPFLP---EDQPMYALQSPILRDPT-----RVIGSLDELAAEYLQRIVDLH-PEGPYQLAGWSVGGNLAL   80
Pys-Pent_TE                20  LAYLP---QDQPLYALQSPILLDPE-----RVVGSLEELAREYLQRILALQ-PQGPYQLAGWSVGGNLAL   80
EntF_TE                    34  SRYLD---PQWSIIGIQSPRPNGPM-----QTAANLDEVCEAHLATLLEQQ-PHGPYYLLGYSLGGTLAQ   94
AB_TE                      32  SRHLN---PNRTLRAIQSPGLIEAD-----AAEVAIEEMATLYIAEMQKMQ-PQGPYFLGGWCFGGAIAY   92
AB_TE.pdb_chainA_s002      32  SRHLN---PNRTLRAIQSPGLIEAD-----AAEVAIEEMATLYIAEMQKMQ-PQGPYFLGGWCFGGAIAY   92
SwrW_TE                    20  ANYLE---KDFNVLGINNYNLFQQ------PHITSLRELAAYYLGHMRLS--PARPVRLLGWSLGGLIAL   78
5UGZ_                      30  RSVLS---DNITLRPLEPAGRGTRIRQ---PLCLTMVDAVADLYQQFVKHY-TGGDYAIFGHSLGGIMAF   92
Consensus_aa:                  ...h........h.shp.................s...hh..hh..l......p.sh.lhG@thGt.hAh
Consensus_ss:                  hhh         eeeee                 hhhhhhhhhhhhhhh        eeeeee hhhhhh
```

**Supplemental figure 6: Structure-based sequence alignment of thioesterase domains from NRPS and PKS proteins.** Green box shows alignment of "QCN" residues of SulTE domain with other TE domain sequences. Presence of "QCN" residues was only found in SulTE sequence. PROMALS3D server was used for alignment. (http://prodata.swmed.edu/promals3d/results/phpONqyVB.result.html)

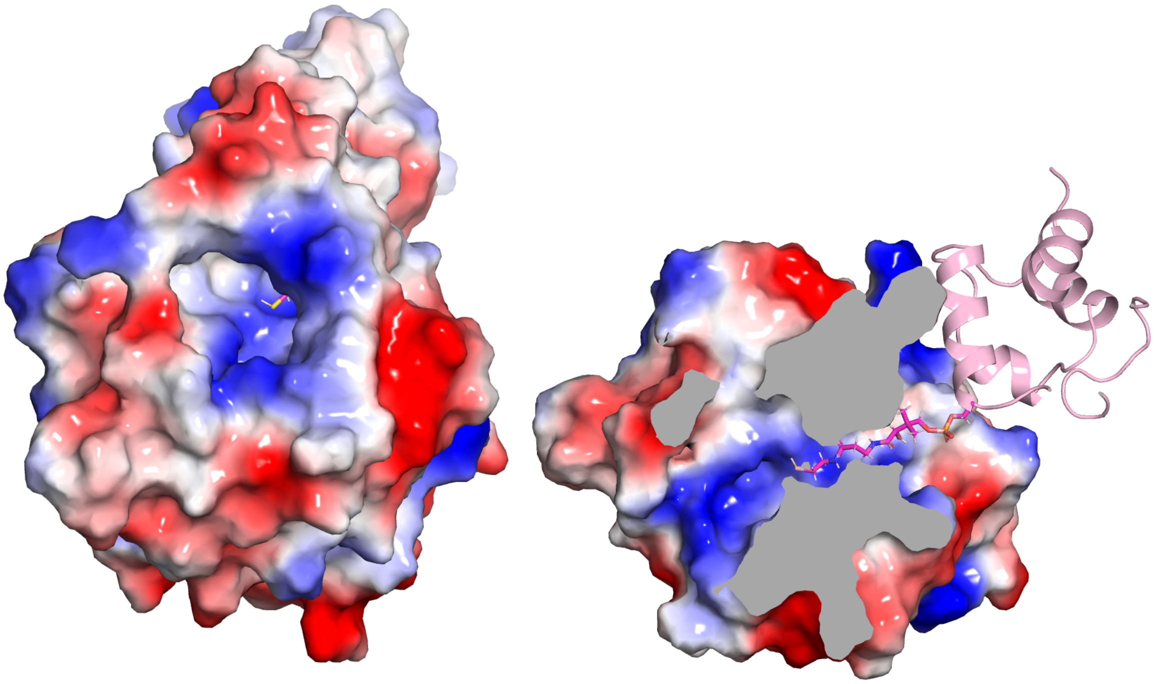

**Supplemental figure 7: Electrostatic potential of SulTE and SulM_PCP-TE structure**. Positively charged substrate binding pocket at the center in SulM_PCP-TE (left) which likely favors binding with negatively charged tripeptide. The phosphopantetheine arm approaches positively charged substrate binding pocket through a tunnel in the SulTE domain (right)

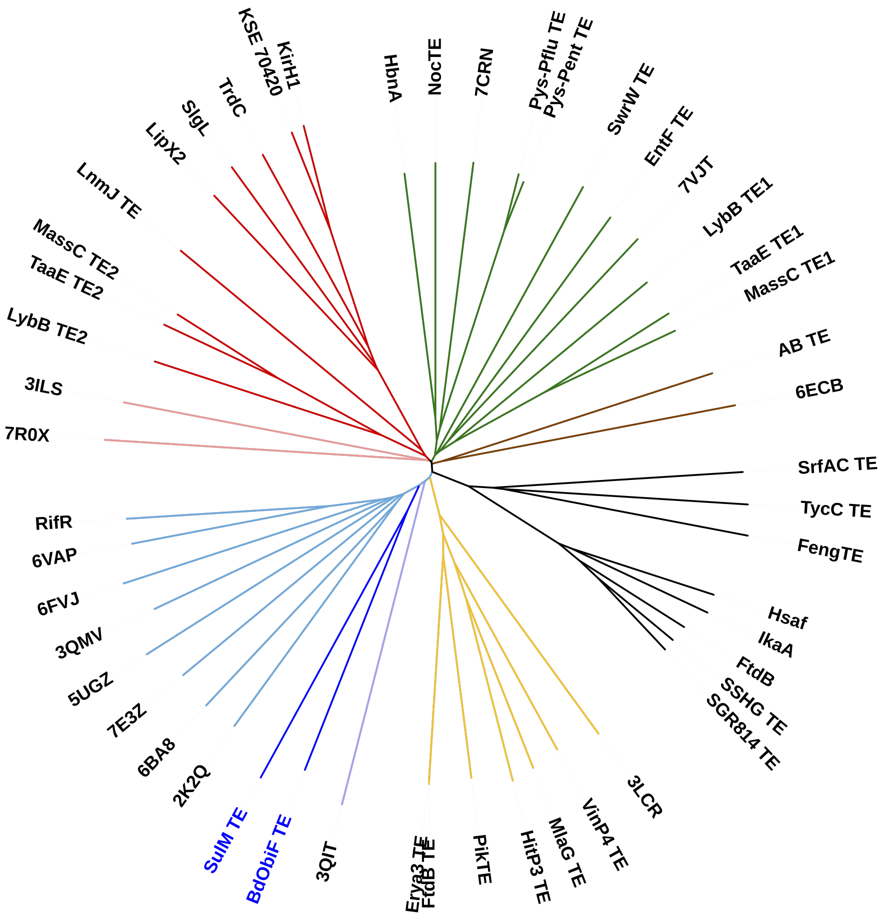

**Supplemental figure 8: Phylogenetic tree of TE domain sequences from NRPS and PKS clusters**. The Interactive Tree of Life software was used to create a phylogenetic tree of thioesterase domains.

```
AQX14441.1_Agrobacterium          ---------------------LHLLSAEEDVGRILVPVAGDLSSAKVRIVAVSNSGGDPI   39
SulM_TE                           ---------------------MEHLEHASEESSSIVLMAGDPATAKAVVVCVANAAGGPV   39
WP_081284758.1_Chromobacterium    -------------------------------QKTLTPIAGDPVQAEAMLLCIANSGGGPV   29
WP_088511513.1_Pseudomonas        ----------------------------STERRVLTRICGDAADADAVFVCIANSGGGPV   32
WP_056247584.1_Flavobacterium     VSLSDSYTVPDLYRNPTIRELAKHIKENKKSNDLLTKMLDVKEKAEVTIIGIPNSAGDPL   60
WP_123275170.1_Chryseobacterium   -------------------------SINSDSVNLLEPMF-VSNSSNETLILVPNSAGEPI   34
                                                                    :  :      :.  .: : *:.* *:

AQX14441.1_Agrobacterium          SFLDLGKAIGRKTEHVAFHAVKLPRIGTTDDAGFREEITRLTNKVAEELSLNANLPTIIY   99
SulM_TE                           NFVDMSRAMPEQASDVAMFGVKLPRTEVDSDGAMLEEVRRLSNAVCDDLLAATDLPAIVF   99
WP_081284758.1_Chromobacterium    SFIEMGRAFSAAGVPLAACAVNLPRNEVDDDAAMVAEVERLAAEVCDELGRMSSLPLMVF   89
WP_088511513.1_Pseudomonas        SFIETGRELAARNGRLAAYAVNLPRNEIDDDAMMVGEVERLTAEVCDSLAQETALPIVVF   92
WP_056247584.1_Flavobacterium     SFSKTVEFIEKDSTNINFYGLKLPRTDPEEGQDMLDMLYKLSEDVVREIQEKVHTPIIIF   120
WP_123275170.1_Chryseobacterium   NFRDLTNSIEAISKEINIYCVKFPRTPLKENENHEKKRKLLLKDLVAEVKEKIRGSITIY   94
                                  .* .  . :      :    :::**    ..         *   :  .:         ::

AQX14441.1_Agrobacterium          GQCNGSALAISLARHLAQETRDVAGLFIGGALMRT--SLSPADNRSDEEILSFLTSLGST   157
SulM_TE                           AQCNGSALALAITRELVRRSADVRALCIGGALMRT--VTGKRDTRTDDEILAFLGKAGST   157
WP_081284758.1_Chromobacterium    AQCNGSALALGVVREWKRRGADLRALCIGGALLRT--EPTPKDVRSDEQIVAFLGSIGST   147
WP_088511513.1_Pseudomonas        AQCNGSALAISIARELTRRGADLRALCIGGALLRT--EHTRKDTRSDETIVDFLRSLGAT   150
WP_056247584.1_Flavobacterium     GQCNGNVLAISIAEILQRINFNVEAICIGAIFPRTRKSLG-FGKRNNSEVVKFLTSLGST   179
WP_123275170.1_Chryseobacterium   GQCNGTALAIGLADELRNNDINIECLYLGALFPLEMKNLVENDKRTDDHILNMLGELGGV   154
                                  .****..**:.:.    .   ::  : :*. :          . *.:. :: :* . *..

AQX14441.1_Agrobacterium          IPSDPAEAAFFLHDFRYDCGFANAYYNQLIE-EVRGHTLEPLNIPLTCMVGTNDEMVRGY   216
SulM_TE                           LPAQPDEQAFFLHDFRYDGWLADVYYNHLVD-LMSRGALEVVDIPVWCLVGSEDPLVPNY   216
WP_081284758.1_Chromobacterium    LPAEADERAFFLHDFSYDCHMADSYYNHLVR-ERAAEAGERIGAPVFCLVGTEDPIVSGY   206
WP_088511513.1_Pseudomonas        LPTRPDEFEFFMQDFRYDCSMADAYYNHLLA-EIDGGRVARIGAPIFNLVGTLDAIVPNY   209
WP_056247584.1_Flavobacterium     IPTDPLDQEFFIKNFRYDSDLAVAGFNHYLEATKKRNRFDKFRMPLHFITGSIDPITKGY   239
WP_123275170.1_Chryseobacterium   FPTESPYKEFFLENIRYDSMMAQTGFHYFYH-QIKEKKFKKFDFPIHSVVGSEDTVTKYY   213
                                  :*:      **:.:: **  :*   ::              .  *:  :.*: * :.  *

AQX14441.1_Agrobacterium          QTDYRLWGKISQEIRLVEYHGFGHYLLRDCPELIADTMLEACSNFCHLESLAS-----   269
SulM_TE                           PVRFQDWSHIGRPVQLVEYAGIGHYLLRDCPEAIARAVGSVWEHVSCKGVTA------   268
WP_081284758.1_Chromobacterium    AERYRDWGLLSDEVSLIEYPGIGHYLLRDCPAELAASLADIWRRVGGK----------   254
WP_088511513.1_Pseudomonas        RARHRDWTHLSDDVQLVEYPGVGHYLLRDCPDLVADTLCSVWKDVRNEG---------   258
WP_056247584.1_Flavobacterium     KRKHKRWHLFAETVDISVIENHGHYLLRDAPAELGNILVNITNKNRKTVNIEYENENK   297
WP_123275170.1_Chryseobacterium   SFRYRKLKKYSNKVSLHVIKGVGHYLHRDAYTQLANILIKMRKK--------------   257
                                     .:     .  : :    . **** **.   :.  : .
```

**Supplemental figure 9: Sequence alignment with SulTE domain homologous proteins**. SulTE protein sequence was used as query for protein BLAST and sequences showing less than 70% sequence similarity were selected. Further screening was done by using structural features of SulTE domain. Sequences were selected by identifying presence of 1) "QCN" residues (highlighted yellow) at the active site, 2) two out of three Arginine residues (highlighted blue) for interacting with substrate residues and 3) catalytic residue Aspartate on position II (highlighted red).

**SulM protein from *Paraburkholderia acidicola***

NCBI Reference Sequence: AOZ21320.1

**SulM Thioesterase Construct**

```
                                    2723
        GSSHHHHHHS SGLVPRGSHM ASEESSSIVL MAGDPATAKA VVVCVANAAG GPVNFVDMSR
2758    AMPEQASDVA MFGVKLPRTE VDSDGAMLEE VRRLSNAVCD DLLAATDLPA IVFAQCNGSA
2818    LALAITRELV RRSADVRALC IGGALMRTVT GKRDTRTDDE ILAFLGKAGS TLPAQPDEQA
2878    FFLHDFRYDG WLADVYYNHL VDLMSRGALE VVDIPVWCLV GSEDPLVPNY PVRFQDWSHI
2938    GRPVQLVEYA GIGHYLLRDC PEAIARAVGS VWEHVSCKGV TA*
```

**SulM PCP-Thioesterase Construct**

```
                                    2660
        GSSHHHHHHS SGLVPRGSHM ERLTEIFRGV LGHAAFGIRD DFFDLGGDSF KAIRIAAKYG
2700    PPLEVTDIYD HPTIEALAEH LEHASEESSS IVLMAGDPAT AKAVVVCVAN AAGGPVNFVD
2760    MSRAMPEQAS DVAMFGVKLP RTEVDSDGAM LEEVRRLSNA VCDDLLAATD LPAIVFAQAN
2820    GSALALAITR ELVRRSADVR ALCIGGALMR TVTGKRDTRT DDEILAFLGK AGSTLPAQPD
2880    EQAFFLHDFR YDGWLADVYY NHLVDLMSRG ALEVVDIPVW CLVGSEDPLV PNYPVRFQDW
2940    SHIGRPVQLV EYAGIGHYLL RDCPEAIARA VGSVWEHVSC KGVTA*
```

**Supplemental figure 10: Protein sequences of SulM thioesterase and PCP-thioesterase constructs**. The twenty residues in red represent the N-terminal $His_6$ tag and thrombin cleavage site, which remained in place for the crystallization experiment. The thioesterase domain begins at residue Ser2728. The thioesterase domain construct begins at Ala2723, containing several residues from the linker following C-terminal α-helix of the PCP; Ala2984 is the natural C-terminus of the SulM protein. The catalytic triad is highlighted in yellow. The PCP-Thioesterase construct constrains the PCP domain (blue), Glu2660 to Ala2723, followed by the thioesterase domain. The GGDS pantetheine binding motif of the PCP is underlined in blue. The catalytic triad is highlighted in yellow, with the mutation cysteine to alanine highlighted in cyan.

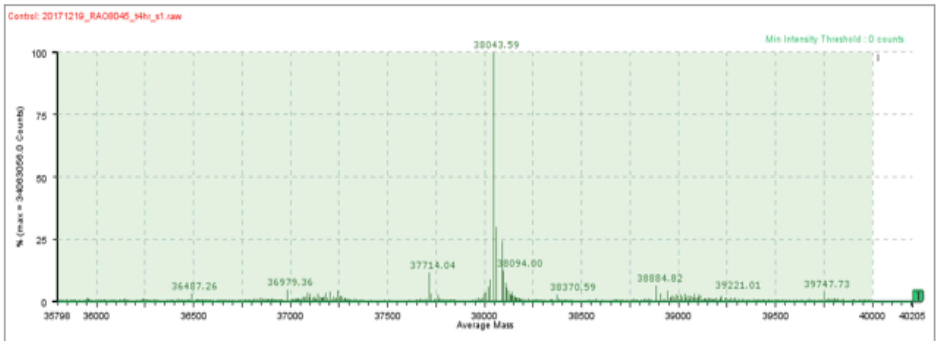

**Supplementary Figure 11. Loading of sulfazecin tripeptide mimic to PCP3-TE C2818A**.
Deconvoluted HRMS spectrum confirmed successful stable loading of tripeptide mimic to the $PCP_3$-TE* C2818A didomain. Calc. 38042 Da (-Met), found 38043 Da.

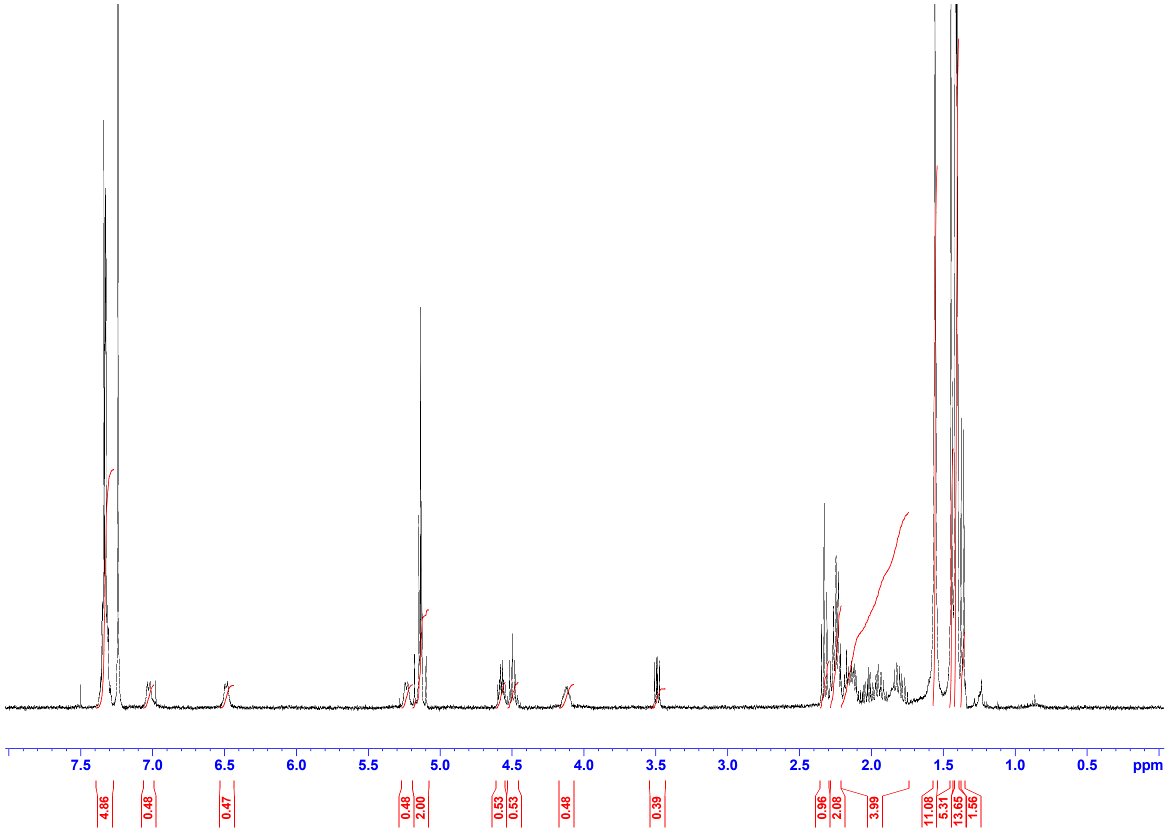

**Supplementary Figure 12. $^1$HNMR spectrum of Compound 3.**

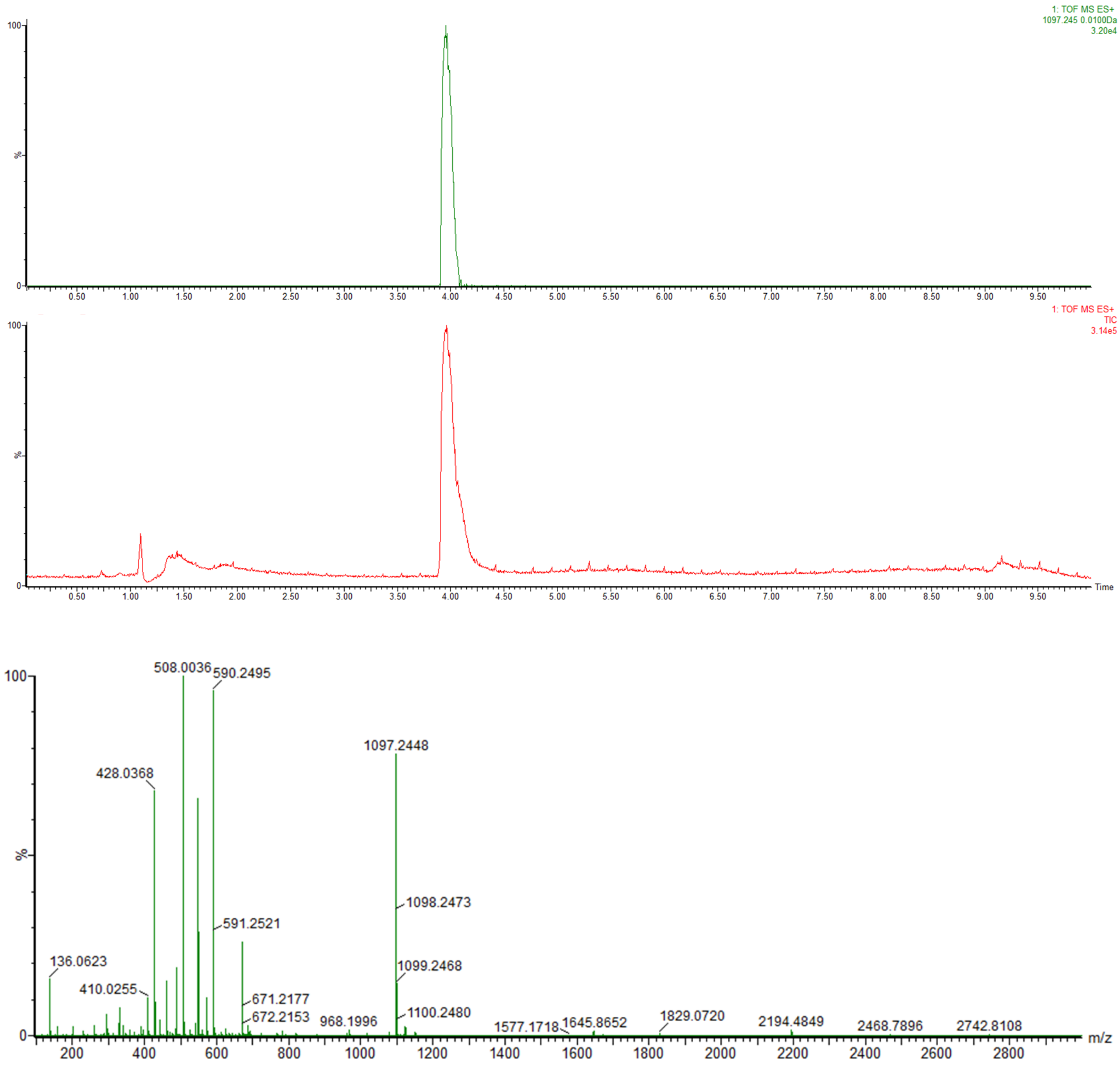

**Supplementary Figure 13. Mass spectrometry of tripeptide-CoA thioester**. Extracted ion chromatogram and total ion chromatogram of **4**. Positive mode ESI: $m/z$ = 1097. 245 ± 0.01. Lower panel shows ESI (+) mass spectrum of **4**.

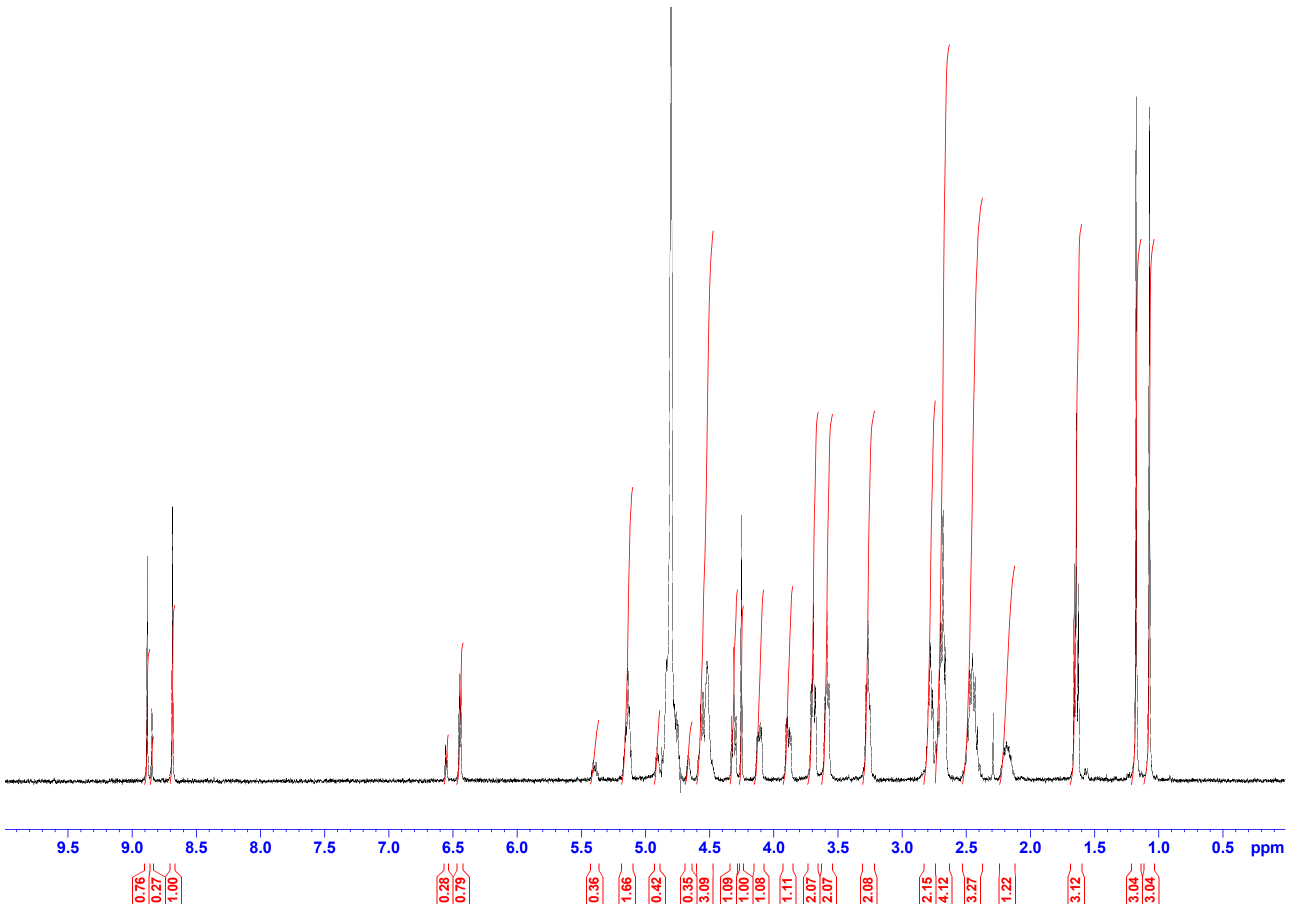

**Supplementary Figure 14. 1HNMR spectrum of Compound 4.**

### Supplemental Tables

**Table S1. Secondary structure of lid regions of structurally characterized NRPS thioesterase domains**.

| Protein | PDB | β6-β7 Loop | # AA | β7-β8 loop details | # AA |
|---|---|---|---|---|---|
| SulM | **8W2C** | Gly2844 – Val2918 | 75 | Gly2923 – Val2946 | 24 |
| | | αL1 Asp2860 – Ala2870 | 11 | α Tyr2932 – Ile2942 | 11 |
| | | αL2 Pro2878 – Ser2907 | 30 | | |
| SrfA-C | **2VSQ** | Val1146 – Asp1211 | 66 | Thr1216 – Arg 1236 | 21 |
| | **1JMK** | αL1a Thr1161 – Asn1171 | 11 | No α helices. | |
| | | αL1b Glu1177 – Ser1181 | 17 | | |
| | | αL2 Lys1185 – Asn1201 | | | |
| Vlm2 | **6ECE** | Leu2489 – Ile2589 | 101 | Thr2594 – 2615 | 22 |
| | | αL1a Thr2506 – Met2517 | 12 | α Ala2598 – Ala2612 | 15 |
| | | αL1b Tyr2535 – Asn2548 | 14 | | |
| | | αL1c Pro2550 – Phe2555 | 6 | | |
| | | αL2 Glu2558 – Met2573 | 16 | | |
| EntF | **5JA1** | Leu1164 – Phe1232 | 69 | Ala1241 – Ile1259 | 19 |
| | | αL1a Pro1169 – Trp1174 | 6 | α Ser1250 – Ser1256 | 7 |
| | | αL1b Glu1185 – Ala1200 | 16 | | |
| | | αL2 Thr1207 – Thr1225 | 19 | | |
| NocB | **6OJD** | Val1805 – Val1865 | 61 | Val1869 – Leu1892 | 24 |
| | | αL1a Asn1809 – Ala1813 | 5 | α Asp1877 – Arg1882 | 6 |
| | | αL1b Asp1818 – Leu1830 | 13 | | |
| | | αL2 Ser1836 – Tyr1856 | 21 | | |
| ObiF | **6N8E** | Gly1172 – Leu1246 | 75 | Gly1251 – Val1274 | 24 |
| | | αL1a Asp1179 – Gly1188 | 10 | α Asp1259 – Ser1271 | 12 |
| | | αL1b Ser1191 – Ile1202 | 12 | | |
| | | αL2 Asp1210 – Glu1235 | 26 | | |
| FenB | **2CB9** | Val1149 – Asn1202 | 54 | Glu1206 – Tyr1231 | 26 |
| | | αL2 Pro1172 – Leu1193 | 22 | α Ser1215 – Leu1220 | 6 |
| | | | | α Trp1223 – Ala1226 | 4 |
| SkyXY | **7CRN** | Leu3753 – Val3844 | 92 | Ala3849 – Ile3875 | 24 |
| | | αL1a Ala3764 – Gln3758 | 5 | α Pro3856 – Val3871 | 16 |
| | | αL1b Asp3770 – Val3782 | 13 | | |
| | | αL1c Asp3795 – Arg3806 | 12 | | |
| | | αL2 Asp3814 – Phe3832 | 19 | | |
| AB3403 | **4ZXI** | Ile1161 – Ile1260 | 100 | Ala1265 – Leu1286 | 22 |
| | | αL1a Ile1167 – Val1171 | 6 | α Glu1277– Leu1281 | 5 |
| | | αL1b Asp1176 – Gly1193 | 18 | | |
| | | αL1c Pro1199 – Glu1205 | 7 | | |
| | | αL1d Ser1207 – Ala1221 | 15 | | |
| | | αL2 Asp1230 – Thr1250 | 21 | | |

**Table S2: Crystallographic Diffraction and Refinement Data**.

| **Data collection** | SulM Thioesterase | SulM PCP-Thioesterase |
|---|---|---|
| **NCBI Accession Code** | WP_096724622.1 | WP_096724622.1 |
| **Residues** | 2778 – 2984 | 2660 – 2940 |
| **PDB** | **8W2C** | **8W2D** |
| **Beamline** | SSRL BL 9-2 | SSRL BL 9-2 |
| **Wavelength (Å)** | 0.979 Å | 0.979 Å |
| **Resolution range (Å)**[a] | 50.0 – 1.9 (2.0 – 1.9) | 47.8 – 2.7 (2.84 – 2.70) |
| **Space group** | $P\ 2_1$ | $P\ 2_1$ |
| **a, b, c (Å)** | 47.09, 83.20, 69.36 | 60.1, 79.1, 70.5 |
| **α, β, γ (°)** | 90.0, 101.4, 90.0 | 90.0, 93.5, 90.0 |
| **Total reflections** | 276587 | 72068 |
| **Unique reflections** | 39586 | 17876 |
| **Multiplicity** | 7.0 (7.0) | 4.0 (3.9) |
| **Completeness (%)** | 95.8 (95.8) | 97.7 (94.5) |
| **Mean I/sigma(I)** | 11.4 (1.5) | 8.1 (1.4) |
| **$R_{merge}$ (%)** | 16.5 (223.9) | 13.9 (94.9) |
| **$R_{pim}$ (%)** | 6.7 (90.3) | 7.7 (53.2) |
| **$CC_{1/2}$** | 0.99 (0.56) | 0.99 (0.65) |
| **Refinement** | | |
| **Resolution range (Å)** | 42.26 – 1.90 (2.00 – 1.95) | 40.46 – 2.70 (2.77 – 2.70) |
| **Reflections, refinement** | 39525 | 17786 |
| **Reflections, $R_{free}$** | 1995 | 1768 |
| **$R_{work}$ (%)** | 17.47 (31.08) | 22.08 (28.55) |
| **$R_{free}$ (%)** | 21.49 (38.77) | 25.69 (33.74) |
| **Protein residues** | 507 | 642 |
| **Ligands of interest** | -- | 2 (Pantetheine) |
| **Water molecules** | 320 | 28 |
| **RMS (bonds) (Å)** | 0.009 | 0.003 |
| **RMS (angles) (°)** | 0.915 | 0.801 |
| **Ramachandran favored (%)** | 98.81 | 97.00 |
| **Ramachandran allowed (%)** | 1.19 | 3.00 |
| **Ramachandran outliers (%)** | 0.0 | 0.0 |
| **Rotamer outliers (%)** | 0.25 | 0.0 |

[a]Values in parentheses are for high resolution shell

**Table S3. Potential uncharacterized β-lactam producing BGCs**. Three BGCs identified from anti-SMASH that may produce uncharacterized β-lactam products. The Stachelhaus code residues in the adenylation domain of the terminal module with high similarity to the sulfazecin cluster are also included.

| Cluster / bacteria | Anti-SMASH predicted product | Stachelhaus code* |
|---|---|---|
| Sulfazecin / Paraburkholderia acidicola | γ-Glu-Ala-DAP | DVWEMNADDK |
| Flavobacterium sp. Leaf82<br>WP_056247584.1<br>NZ_LMMA01000010.1 | Orn-Gly-Asp-Asp-Xxx-Thr-DAP | DIWEINTDDK |
| *Agrobacterium tumefaciens*<br>BGC0001671<br>KY452017.1 | Xxx-DAP | DIWEITADDK |
| Chryseobacterium sp. G0240<br>WP_123275170.1<br>NZ_RJTV01000006.1 | Asp-Leu-D-Orn-DAP | DIWEVNTDDK |

* Stachelhaus code residues in adenylation domain of the last modules
